# Species-Specific Roles of RIPK1 and TRADD in TNF-Induced Cell Death Reveal a Translational Gap Between Mouse Models and Human Biology

**DOI:** 10.64898/2026.06.28.735126

**Authors:** Zhao Deng, Buhao Deng, Jingyi Wang, Jiafan Yuan, Kai Yu, Yan Liu, Haiwen Lin, Youwei Ai, Bo Yan

**Affiliations:** Institute of Genetics and Developmental Biology, Chinese Academy of Sciences, Beijing 100101, China; University of Chinese Academy of Sciences, Beijing 100101, China; Tianjin Neurological Institute, Tianjin Medical University General Hospital, Tianjin 300052, China; College of Life Science, Yunnan University, Kunming 650091, China

## Abstract

Mouse models have historically been central to studies of TNF-induced cell death and guided pharmaceutical translation into clinic, based on the assumption that TNF signaling is conserved between human and mouse. Here, our work uncovers critical species-specific differences between the two. By systematically dissecting the roles of RIPK1, TRADD, and sensitivity to RIPK1 inhibitors in TNF signaling—including RIPK1 kinase-dependent and-independent apoptosis—we found that both apoptosis modalities diverge between human and mouse cells. In mouse cells, RIPK1 suppresses TRADD-mediated kinase-independent apoptosis, whereas in human cells, RIPK1 and TRADD act redundantly. Moreover, RIPK1 inhibitors block kinase-dependent apoptosis in mouse but not human cells, despite effectively inhibiting RIPK1 S166 phosphorylation. Cross-species complementation revealed that these discrepancies stem not from RIPK1 itself but from cell-context differences. These findings echo the clinical failures of RIPK1 inhibitors despite efficacy in mouse models and underscore the need for humanized models and therapeutics that more faithfully predict clinical outcomes.

## Introduction

Tumor necrosis factor α (TNF) is a pleiotropic cytokine whose upregulation exacerbates tissue damage in various acute and chronic inflammatory and autoimmune diseases. These include rheumatoid arthritis, psoriasis, ankylosing spondylitis, inflammatory bowel diseases (IBD, including Crohn’s disease and ulcerative colitis), and other inflammatory disorders affecting both the peripheral and central nervous systems.^1,2^ Since their clinical introduction in 1998, TNF-targeting biologics—particularly TNF-neutralizing antibodies—have become some of the most successful drugs in history, generating over $40 billion in global annual sales. However, approximately one-third of patients discontinue these biologics due to lack of efficacy or adverse effects.^2,3^ Thus, developing novel therapeutic strategies targeting the TNF pathway for inflammatory diseases has long been an active field of research.

TNF signaling is highly complex. Under basal conditions, TNF primarily activates NF-κB and MAPK pathways, promoting cell survival.^4,5^ However, when cell death checkpoints are compromised, TNF can trigger various forms of programmed cell death, including RIPK1 kinase-independent apoptosis, RIPK1 kinase-dependent apoptosis, RIPK1 kinase-dependent necroptosis, and other less characterized modalities.^6–9^

Historically, mouse cells and models have been indispensable for mechanistic studies of TNF signaling and preclinical research. Conclusions are often drawn indiscriminately from both human and mouse studies under the assumption that signaling pathways are conserved across species, and these are then presented to the broader TNF community and even non-specialists.^6,10–19^ Typically, TNF signaling is introduced in the following manner. TNF binding to its transmembrane receptor TNFR1 induces trimerization of the receptor’s intracellular death domain (DD).^20^ This facilitates the recruitment of two additional DD-containing proteins, RIPK1 and TRADD, to form the membrane-bound complex I via homotypic DD interactions. Within complex I, TRADD recruits E3 ligases such as TRAF2/5, cIAP1/2, and the LUBAC complex (HOIL1, HOIP, and SHARPIN).^16,17,21–24^ These ligases catalyze polyubiquitination of RIPK1 and potentially other substrates, enabling downstream recruitment of kinases such as IKKα/β and TAK1 to activate NF-κB and MAPK signaling, respectively.^19,25–27^

Under certain stress conditions, complex I dissociates from membrane and forms complex II to induce cell death. These conditions include: (i) inhibition of NF-κB target gene expression via transcriptional (e.g., actinomycin D or D-galactosamine) or translational (e.g., cycloheximide, CHX) inhibitors; (ii) co-stimulation with inflammatory cytokines (e.g., IFNγ, TWEAK, or lymphotoxin-β); or (iii) pharmacologic inhibition of cIAP1/2 (e.g., SMAC mimetic, or BV6) or TAK1 (e.g., 5Z-7-oxozeaenol, 5Z7).^6,28–30^ TNF/CHX and TNF/IFNγ co-treatment activates RIPK1 kinase-independent apoptosis, while TNF/SMAC (TNF/BV6) and TNF/5Z7 co-treatment activates RIPK1 kinase-dependent apoptosis.^2,9,28^

Upon dissociation of complex I components such as RIPK1 and TRADD into the cytosol, they recruit FADD and caspase-8 to form complex II, initiating apoptosis. Caspase-8 inhibition can prevent apoptosis in some cell types, allowing survival; however, in cells expressing RIPK3 and MLKL—such as the human HaCaT, HT29, U937 cells, macrophages, and intestinal organoids, and mouse iMEF, L929 cells, macrophages, and intestinal organoids used in this study—caspase-8 inhibition redirects signaling toward RIPK1 kinase-dependent necroptosis.^31–36^

While TNF-induced NF-κB and MAPK activation is largely inflammatory, excessive apoptosis or necroptosis—especially when surpassing immune cell clearance—can cause tissue damage.^37^ TNF-driven cell death is increasingly implicated in inflammation-related pathologies. Consequently, small-molecule RIPK1 kinase inhibitors have been actively pursued to treat TNF-related diseases, particularly in patients unresponsive to TNF biologics and in central nervous system (CNS) inflammation.^3^ Despite promising efficacy in murine models and the successful development of stable, blood-brain barrier (BBB)-permeable RIPK1 inhibitors,^3,38^ none have yet succeeded in clinical trials.^39–42^ This failure may partly stem from the widely accepted but not systematically verified assumption that TNF signaling is conserved between human and mouse cells.

Although individual knockouts of RIPK1 and TRADD have been investigated in certain aspects of TNF signaling, and the effects of RIPK1 inhibitors—particularly in necroptosis—have been repeatedly tested,^43–54^ no comprehensive or species-specific comparative studies have been conducted to date. In this study, we systematically compare the roles of RIPK1 and TRADD in TNF-induced NF-κB and MAPK activation, kinase-dependent and-independent apoptosis, necroptosis, and their responsiveness to RIPK1 kinase inhibitors in human and mouse cells. Our findings challenge the assumption of functional equivalency between species, provide insights into the clinical failure of RIPK1 inhibitors, propose new directions for their development, and argue for the use of humanized mouse models in inflammatory disease research.

## Results

### RIPK1 and TRADD are functionally redundant in kinase-independent apoptosis in human cells

To systematically evaluate the roles of the DD-containing proteins RIPK1 and TRADD downstream of TNFR1 signaling in human cells, we individually or simultaneously knocked out RIPK1 and TRADD in a panel of cell lines: HeLa (cervical cancer), MCF-7 (breast cancer), and HaCaT (keratinocyte) cells (Figure S1A), as well as HCT116 (colon cancer) cells (Figure S2A) and Jurkat (T-cell leukemia) cells (Figures S3B and S3E). In HeLa cells, RIPK1 knockout (KO) abrogated RIPK1 kinase-dependent apoptosis—triggered by TNF/SMAC or TNF/BV6, or by TNF/5Z7 in other cell lines—but did not affect RIPK1 kinase-independent apoptosis induced by TNF/CHX or TNF/IFNγ (Figures 1A–1C). Re-expression of RIPK1 in the knockout background cells restored RIPK1 kinase-dependent cell death (Figures S1B and S1C). In contrast, TRADD KO did not impair either form of apoptosis (Figures 1A–1C). However, double knockout (dKO) of RIPK1 and TRADD completely blocked RIPK1 kinase-independent apoptosis (Figures 1B–D), suggesting functional redundancy. Notably, dKO did not impair intrinsic apoptosis triggered by BCL2/MCL1 inhibitors (ABT737/S63845), which served as a control (Figure 1E).

**Figure 1.**
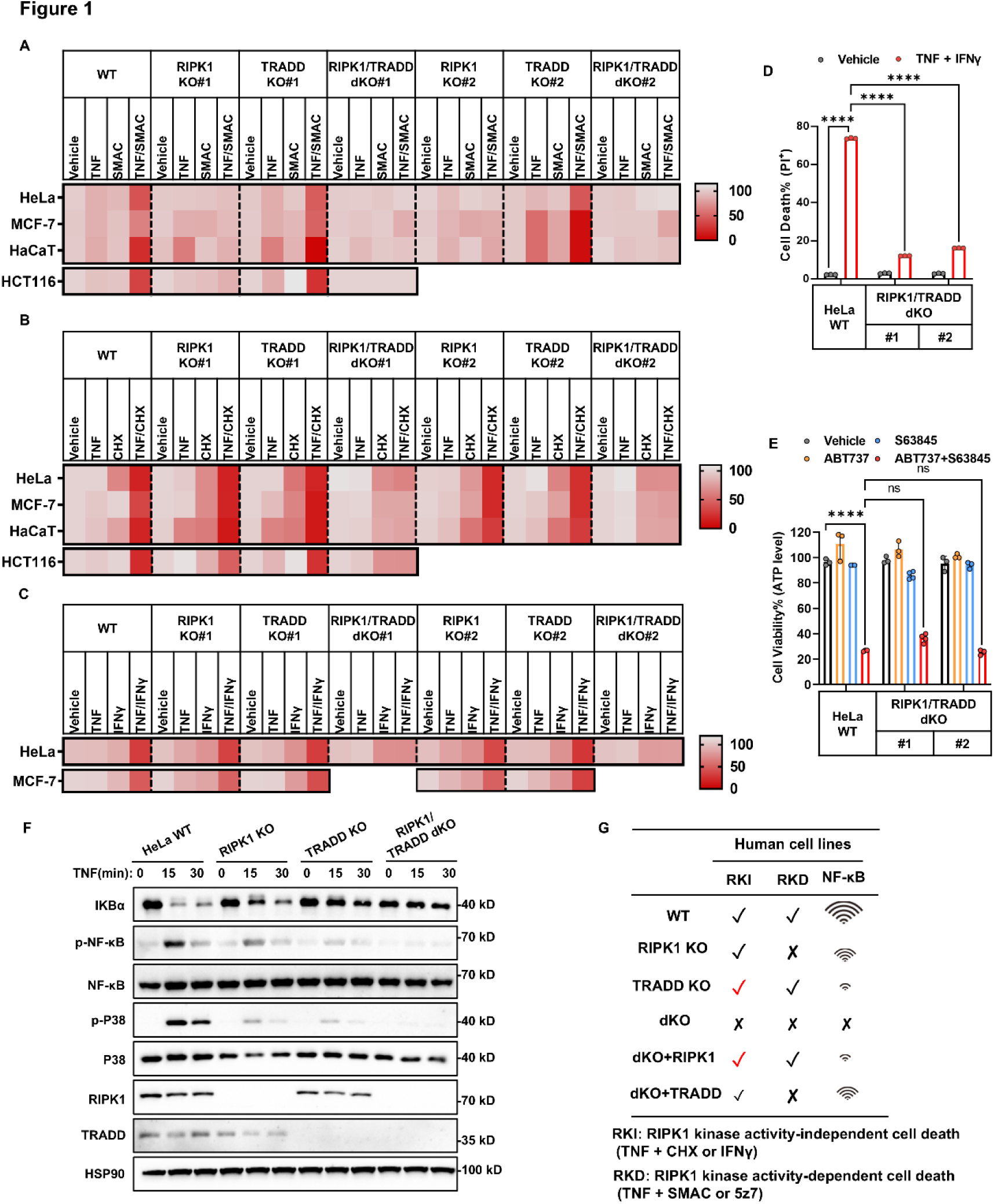
RIPK1 and TRADD are functionally redundant in kinase-independent apoptosis in HeLa cells. (A-C) Heatmaps depict normalized cell viability (%) across human cell lines with the indicated genotypes (WT, RIPK1 KO, TRADD KO, and RIPK1/TRADD dKO) following treatment as indicated. Values represent normalized mean cell viability (%) from three biological replicates. Cell viability was measured by ATP-based luminescence assay. (D) HeLa WT and RIPK1/TRADD dKO clones (#1 and #2, generated with different gRNAs) were treated with TNF/IFNγ for 48 h. Cell death was quantified by propidium iodide (PI) staining followed by flow cytometry (n = 3). (E) HeLa WT and dKO cells were treated with ABT737 and S63845 for 8 h to induce intrinsic apoptosis. Cell viability was measured by ATP-based luminescence assay (n = 3). (F) Immunoblot analysis of NF-κB pathway activation in wild-type (WT), RIPK1 KO, TRADD KO, and RIPK1/TRADD dKO HeLa cells treated with TNF for 0, 15, or 30 minutes. (G) A summary illustrates the genetic requirements of RIPK1 and TRADD in TNF signaling in human cells. The check mark (✓) indicates that the inducer successfully triggers cell death, while a cross mark (✕) indicates that cell death induction by the stimulus is inhibited. The WiFi-like icon represents NF-κB signaling activity, with more signal bars indicating stronger pathway activation. The data in all graphs are showed as mean ± SD. Representative immunoblots from three independent experiments are shown. Statistical significance was determined using two-way ANOVA for group comparisons (D, E): ****P < 0.0001; ns, not significant.

To further confirm this redundancy, we reintroduced either RIPK1 or TRADD into RIPK1/TRADD-dKO HeLa cells. Re-expression of RIPK1 restored both kinase-dependent and-independent apoptosis (Figures S1F-S1I), whereas TRADD alone restored only kinase-independent apoptosis (Figures S1J-S1L). Similar patterns were observed in other human cell types, including HCT116 (Figures 1A, 1B, S2C-S2E, and S2G-S2I), MCF-7 (Figures 1A-1C), HaCaT (Figures 1A and 1B), and Jurkat (Figures S3B-S3G). These results demonstrate that in human cells, RIPK1 and TRADD are functionally redundant for kinase-independent apoptosis, but RIPK1 alone is essential for kinase-dependent apoptosis (Figure 1G).

### RIPK1 and TRADD additively contribute to NF-κB activation in human cells

TRADD has traditionally been viewed as an upstream adaptor that recruits E3 ligases to promote RIPK1 polyubiquitination, thereby enabling NF-κB and MAPK activation.^26,55–57^ This model implies a linear relationship, where TRADD acts upstream of RIPK1. Accordingly, it would be expected that knockout of either TRADD or RIPK1 would yield similar levels of inhibition. However, we found that individual KO of either RIPK1 or TRADD only partially impaired TNF-induced NF-κB and MAPK activation, a pattern consistently observed across multiple cell lines, including HeLa (Figures 1F, S1D, and S1E), HCT116 (Figures S2B and S2F), and HaCaT cells (S3A). In contrast, RIPK1/TRADD double knockout nearly abolished this activation (Figures 1F and S2B).

Interestingly, TRADD KO caused a greater reduction in pathway activation than RIPK1 KO alone (Figures 1F, S2B and S3A), suggesting TRADD plays a more dominant role. These findings support three key conclusions: (i) both RIPK1 and TRADD are independently capable of supporting NF-κB and MAPK signaling; (ii) TRADD contributes more robustly than RIPK1; and (iii) RIPK1 and TRADD function in a parallel and additive—not linear—manner in mediating TNF-induced NF-κB and MAPK activation (Figure 1G).

### RIPK1 and TRADD are functionally different in the assembly of complex I and II

To further investigate the mechanistic basis of the functional differences between RIPK1 and TRADD, we assessed the formation of membrane-bound complex I and cytosolic complex II. Complex I was analyzed by FLAG-TNF immunoprecipitation, and complex II formation was visualized using confocal microscopy in cells expressing FADD-GFP. In wild-type (WT) cells, TNF/SMAC or TNF/CHX treatment induced rapid formation of complex I within 10 minutes, which subsequently dissociated from the membrane at later time points (Figures 2A and 2B). This dissociation is presumed to lead to complex II formation in the cytosol to initiate cell death. The pan-caspase inhibitor z-VAD was used in all conditions to inhibit cell death, and therefore to preserve cellular protein content for immunoprecipitation and to stabilize complex II puncta.

**Figure 2.**
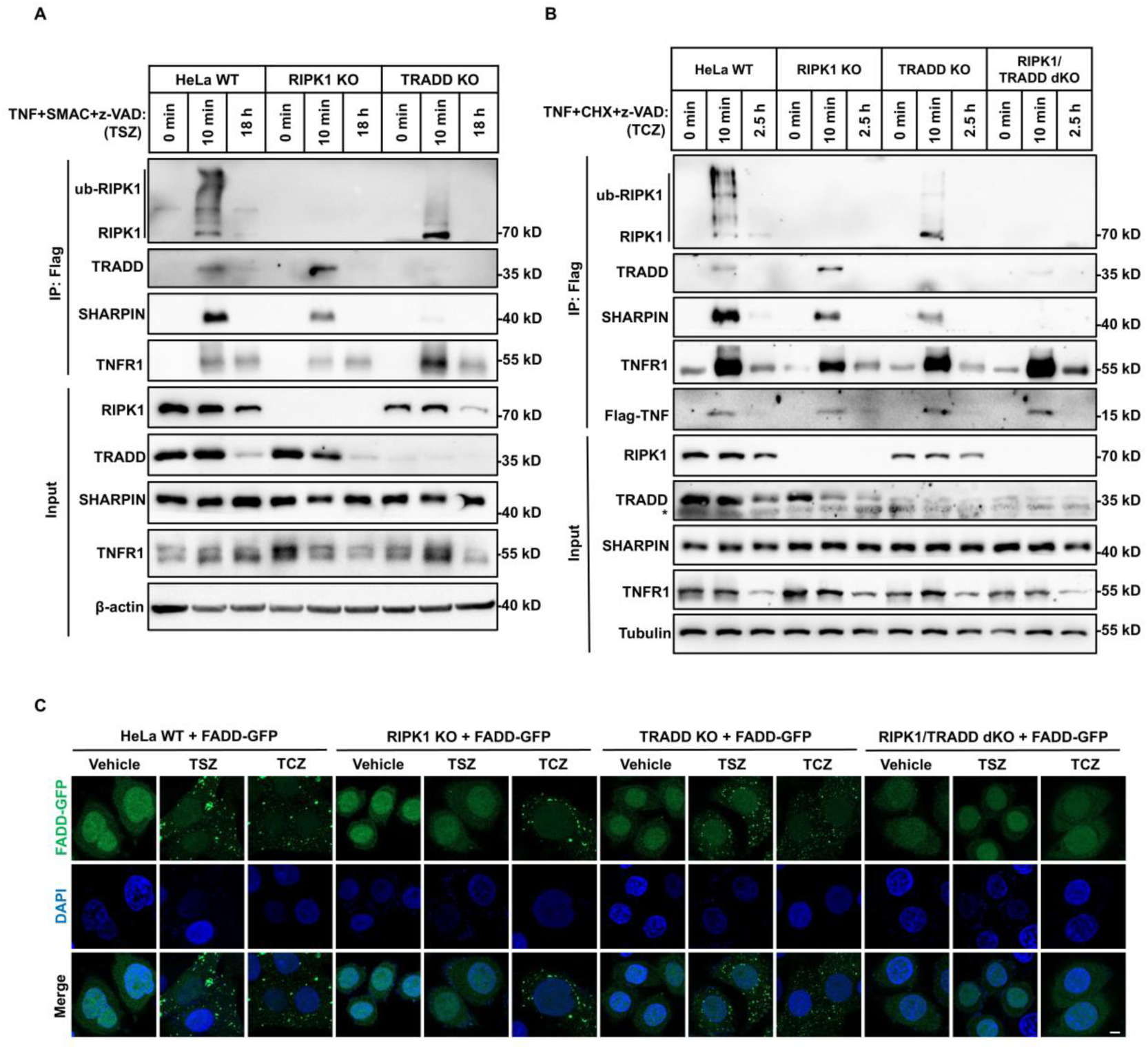
RIPK1 and TRADD differentially regulate complex II assembly in RIPK1 kinase-dependent and-independent apoptosis. (A) HeLa WT, RIPK1 KO, TRADD KO, and RIPK1/TRADD dKO cells were treated with FLAG-TNF, SMAC, and z-VAD (TSZ) for 0, 10 minutes, or 18 hours. Membrane-bound complex I was isolated by immunoprecipitation using anti-FLAG antibody. Immunoblotting was performed to detect proteins as indicated. (B) HeLa WT, RIPK1 KO, TRADD KO, and RIPK1/TRADD dKO cells were treated with FLAG-TNF, CHX, and z-VAD (TCZ) for 0, 10 minutes, or 2.5 hours. Complex I components were isolated by anti-FLAG immunoprecipitation and analyzed by immunoblotting. Asterisk in the immunoblot indicates non-specific band. (C) HeLa WT, RIPK1 KO, TRADD KO, and dKO cells stably expressing FADD-GFP were treated with TSZ (18 h) or TCZ (2.5 h). Cytosolic complex II formation was visualized by confocal microscopy as GFP-positive puncta. Nuclei were counterstained with DAPI. Scale bar, 5 μm. Representative immunoblots and immunofluorescence images from three independent experiments are shown.

In RIPK1 KO cells, recruitment of TRADD to TNFR1 was increased, whereas in TRADD KO cells, levels of ubiquitinated RIPK1 in complex I were reduced, with a corresponding increase in non-ubiquitinated RIPK1 (Figures 2A and 2B). The diminished RIPK1 ubiquitination observed in TRADD KO cells may result from reduced recruitment of E3 ligases, as indicated by decreased levels of the LUBAC subunit SHARPIN. Notably, the reduction in SHARPIN levels in RIPK1 KO, TRADD KO, and dKO cells correlated with the extent of NF-κB and MAPK inhibition observed in each genotype (Figures 2A and 2B vs. 1F).

Both TNF/SMAC/z-VAD and TNF/CHX/z-VAD induced formation of cytosolic complex II, marked by FADD-GFP puncta, in WT, necroptosis deficient HeLa cells (Figure 2C). Although RIPK1 KO cells exhibited comparable complex I formation under both treatments (Figures 2A and 2B), only TNF/CHX/z-VAD—but not TNF/SMAC/z-VAD—induced efficient complex II formation, indicating that RIPK1 is required for kinase-dependent, but not kinase-independent, complex II assembly (Figure 2C). TRADD KO had no effect on complex II formation under either condition, whereas RIPK1/TRADD dKO completely abolished complex II assembly (Figure 2C), consistent with the observed apoptosis phenotypes (Figure 1).

Together, these data suggest that RIPK1 and TRADD cooperatively recruit SHARPIN and E3 ligases to complex I, thereby enabling full NF-κB and MAPK activation. For apoptosis, they are redundant in assembling complex II under kinase-independent conditions, whereas RIPK1 is essential in kinase-dependent settings.

### TRADD but not RIPK1 is indispensable for kinase-independent apoptosis in mouse cells

To determine whether the functional roles of RIPK1 and TRADD are conserved in mice, we performed single and double knockouts in EMT6 (mammary carcinoma), iMEF (immortalized mouse embryonic fibroblasts), and single knockouts in L929 (fibroblast), and CT26 (colon carcinoma) (Figure S4A). In these cell types, RIPK1 KO impaired TNF/SMAC-or TNF/5Z7-induced kinase-dependent apoptosis (Figures 3A and 3B), mirroring human cells. However, unlike in human cells, RIPK1 KO in mouse cells enhanced TNF/CHX-induced kinase-independent apoptosis (Figure 3C). This increase was dependent on TRADD, as the elevated apoptosis in RIPK1 KO cells was abolished in RIPK1/TRADD dKO cells (Figure 3C). These results indicate that in mouse cells, RIPK1 may play an inhibitory role in kinase-independent apoptosis.

**Figure 3.**
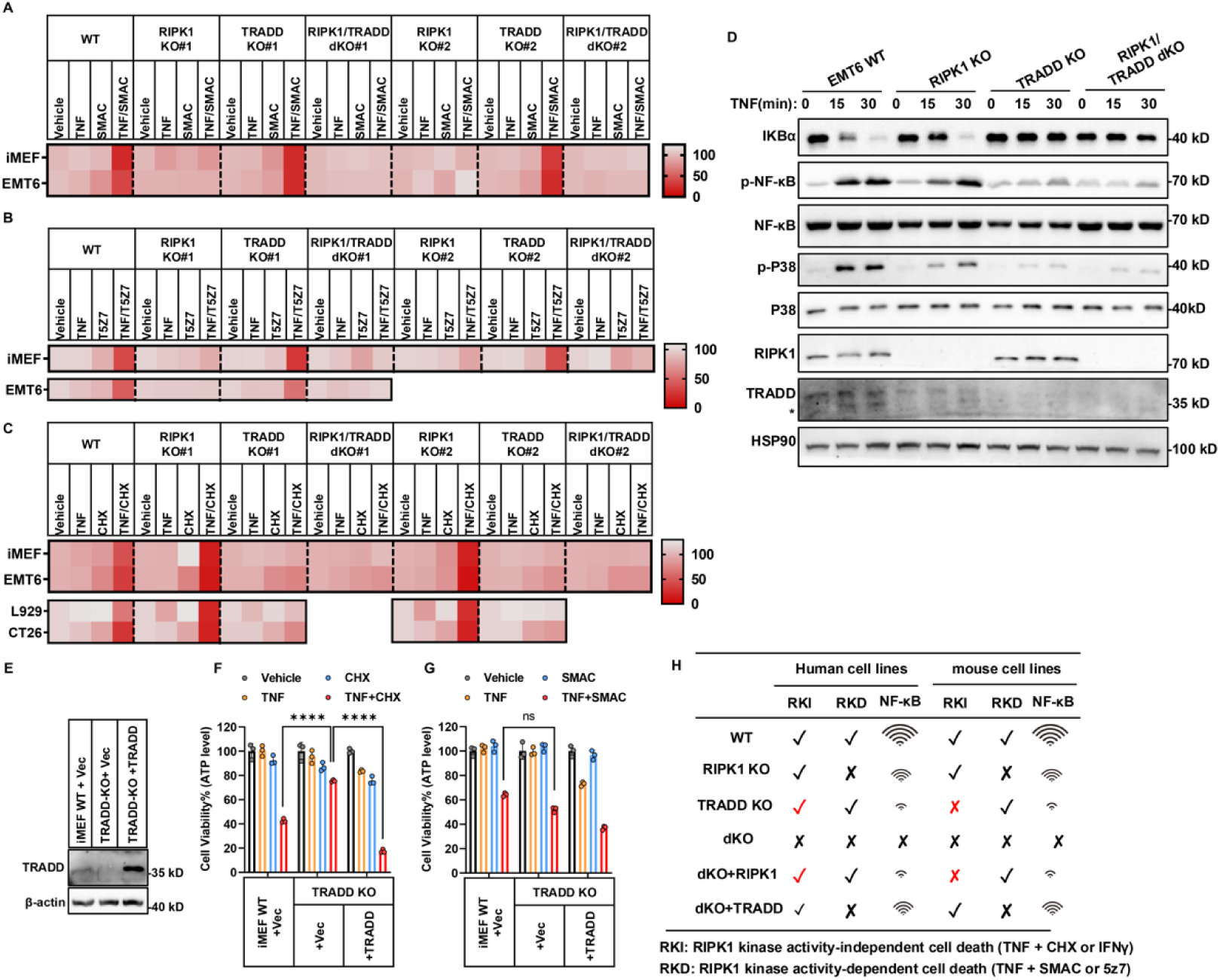
TRADD is indispensable for kinase-independent apoptosis in mouse EMT6 cells. (A-C) Heatmaps showing cell viability across a panel of mouse cell lines with the indicated genotypes (WT, RIPK1 KO, TRADD KO, and RIPK1/TRADD double KO) following treatment as indicated. Values represent normalized mean cell viability (%) from three biological replicates. Cell viability was measured by ATP-based luminescence assay. (D) EMT6 WT, RIPK1 KO, TRADD KO, and RIPK1/TRADD dKO cells were treated with TNF for 0, 15, or 30 minutes. NF-κB and MAPK pathway activation was assessed by immunoblotting. Asterisk in the immunoblot indicates non-specific band. (E-G) iMEF TRADD KO and TRADD-reconstituted cells were assessed. Re-expression of WT TRADD was confirmed by immunoblotting (E). Cells were treated with TNF/CHX for 8 h (F) or TNF/SMAC for 24 h (G), and cell viability was measured by ATP-based luminescence assay (n = 3). (H) A summary illustrates the genetic requirements of RIPK1 and TRADD in TNF signaling in human and mouse cells. A check mark (✓) indicates that the inducer successfully triggers cell death, while a cross mark (✕) indicates that cell death induction by the stimulus is inhibited. The WiFi-like icon represents NF-κB signaling activity, with more signal bars indicating stronger pathway activation. The data in all graphs are showed as mean ± SD. Representative immunoblots from three independent experiments are shown. Statistical significance was determined using two-way ANOVA for group comparisons (F, G): ****P < 0.0001; ns, not significant.

Consistent with human results (Figure 1A), TRADD KO did not affect TNF/SMAC-induced kinase-dependent apoptosis in mouse cells (Figure 3A). However, in contrast to human cells (Figure 1B), TRADD KO completely abolished TNF/CHX-induced kinase-independent apoptosis (Figure 3C). Re-expression of TRADD in KO cells restored sensitivity to RIPK1 kinase-independent apoptosis, both in iMEF (Figures 3E-3G) and EMT6 (Figures S4B-S4D). Similar to human cells, RIPK1/TRADD dKO in mouse cells suppressed both forms of apoptosis (Figures 3A-3C). Interestingly, although RIPK1’s role in TNF/CHX-induced apoptosis differs between human and mouse cells, RIPK1 KO similarly blocked TNF/SMAC/z-VAD or TNF/CHX/z-VAD-induced necroptosis in both human HaCaT cells (Figures S3H and S3I) and mouse cells iMEF cells (Figures S4E and S4F).

The roles of RIPK1 and TRADD in NF-κB and MAPK activation in mouse cells resembled those in human cells (Figures 3D and S4B). Altogether, these data reveal a fundamental species difference: while TRADD is indispensable for kinase-independent apoptosis in mouse cells but not in human cells, RIPK1 is functionally redundant in human cells, but in mouse cells it is dispensable and can even act as an inhibitory factor (Figure 3H).

### TRADD is required for complex II formation during kinase-independent apoptosis in mouse cells

Given the essential role of TRADD in kinase-independent apoptosis in mouse cells, we examined complex I and II formation in TRADD KO mouse cells using iMEF and EMT6 cells expressing FADD-GFP (Figure 4A, 4B, 4E and 4F). TRADD KO cells retained reduced RIPK1 ubiquitination in complex I under both TNF/CHX and TNF/SMAC treatment, indicating loss of E3 ligase recruitment (Figure 4C). However, only TNF/CHX-induced complex II formation was impaired, as indicated by reduced FADD–caspase-8–RIPK1 complex detected by co-immunoprecipitation and diminished FADD-GFP puncta (Figures 4D and 4G). TNF/SMAC-induced complex II formation remained intact (Figures 4D and 4G), reinforcing TRADD’s selective role in kinase-independent cell death in mouse cells.

**Figure 4.**
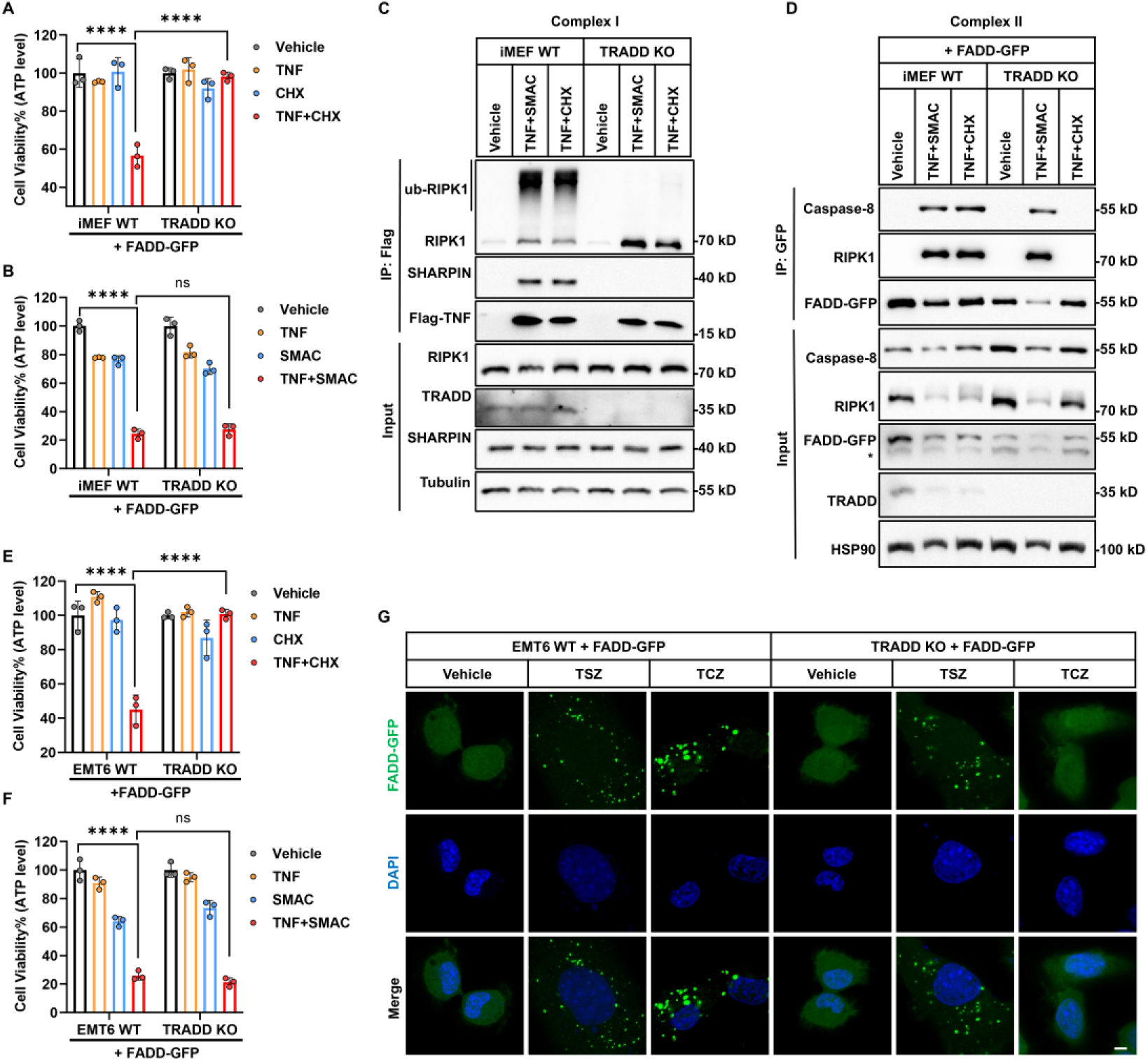
TRADD is essential for complex II assembly in RIPK1 kinase-independent apoptosis in mouse cells. (A, B) iMEF cells stably expressing FADD-GFP (WT or TRADD KO) were treated with TNF/CHX for 8 h (A) or TNF/SMAC for 24 h (B). Cell viability was measured by ATP-based luminescence assay (n = 3). (C) iMEF WT and TRADD KO cells were treated with Flag-TNF/SMAC or Flag-TNF/CHX for 5 min. Membrane-bound complex I was immunoprecipitated using anti-FLAG antibody, and associated proteins were analyzed by immunoblotting. (D) iMEF cells (WT or TRADD KO) stably expressing FADD-GFP were treated with TNF/SMAC (8 h) or TNF/CHX (2 h). Cytosolic complex II was immunoprecipitated using anti-GFP antibody, and co-precipitated proteins were analyzed by immunoblotting. Asterisk in the immunoblot indicates non-specific band. (E, F) EMT6 cells (WT or TRADD KO) stably expressing FADD-GFP were treated with TNF/CHX for 8 h (e) or TNF/SMAC for 12 h (f). Cell viability was measured by ATP-based luminescence assay (n = 3). (G) EMT6 cells (WT or TRADD KO) stably expressing FADD-GFP were treated with TSZ for 6 h or TCZ for 2 h. Representative confocal images show cytosolic FADD-GFP puncta and nuclear DAPI staining. Scale bar, 5 μm. The data in all graphs are showed as mean ± SD. Representative immunoblots and immunofluorescence images from three independent experiments are shown. Statistical significance was determined using two-way ANOVA for group comparisons (A, B, E, F): ****P < 0.0001; ns, not significant.

### Cellular context, not RIPK1 protein identity, governs kinase-independent apoptosis regulation

Knocking out TRADD in mouse cells blocks kinase-independent apoptosis, indicating that the remaining mouse RIPK1 is insufficient to support this pathway—unlike its human counterpart. To clarify whether this species difference is due to intrinsic properties of RIPK1 or the cellular environment, we re-expressed human or mouse RIPK1 in RIPK1/TRADD-dKO HeLa cells. Both human and mouse RIPK1 restored TNF/SMAC-induced kinase-dependent apoptosis (Figures 5A, 5B, and 5D), indicating that both proteins are competent in this context. Unexpectedly, mouse RIPK1 also restored TNF/CHX-induced kinase-independent apoptosis and complex II formation to the same extent as human RIPK1 (Figures 5C and 5D). These results suggest that mouse RIPK1 is functionally competent to mediate kinase-independent apoptosis, but only in the human cellular environment—implying that species-specific cellular factors, not RIPK1 itself, determine this functional divergence.

**Figure 5.**
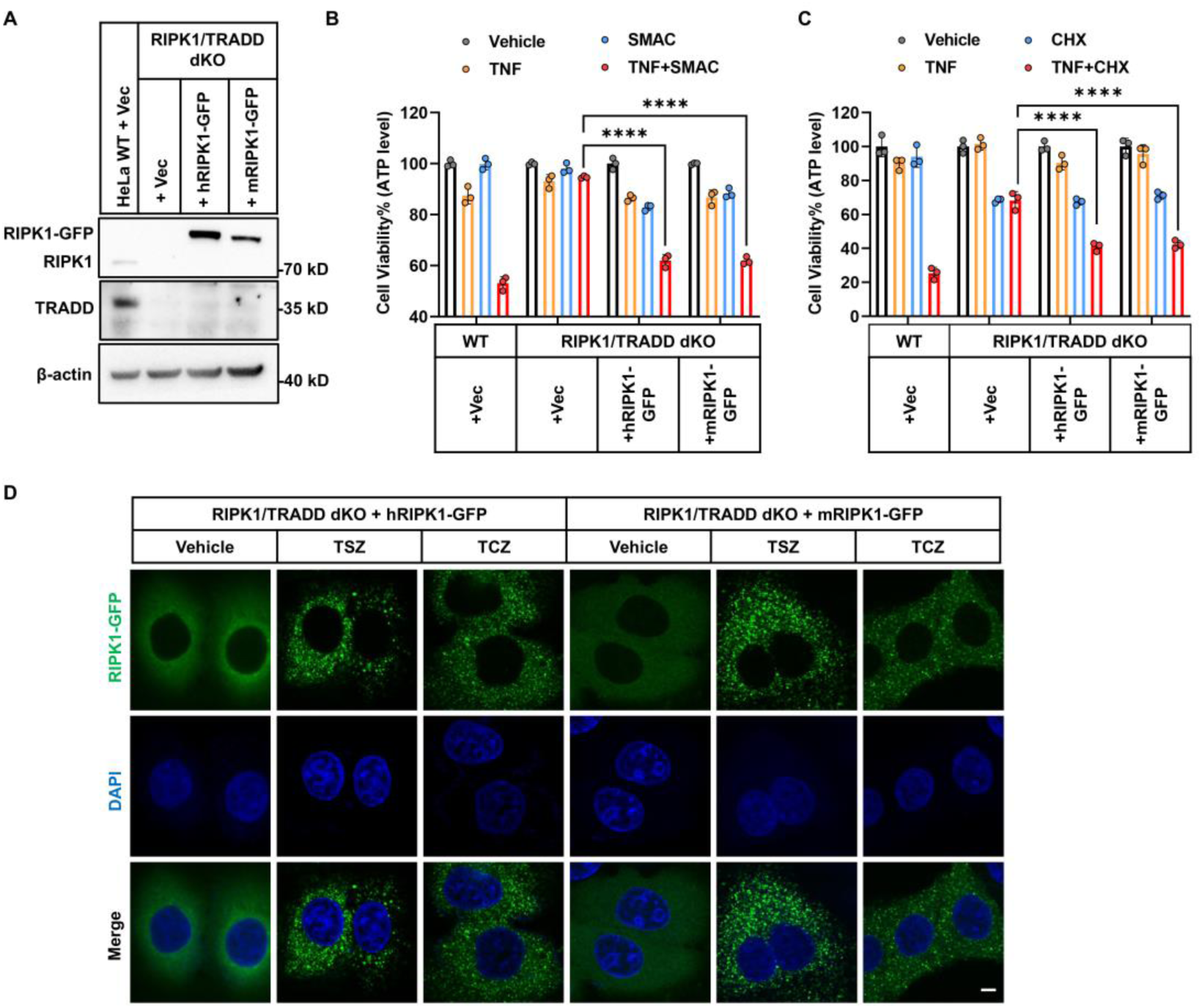
Mouse RIPK1 can mediate RIPK1-independent apoptosis in human HeLa cells. (A-C) HeLa cells (WT, RIPK1/TRADD dKO, dKO reconstituted with human RIPK1-GFP, or dKO reconstituted with mouse RIPK1-GFP) were analyzed. Expression of RIPK1 and TRADD was confirmed by immunoblotting (A). Cells were treated with TNF/SMAC for 48 h (B) or TNF/CHX for 8 h (C), and cell viability was measured by ATP-based luminescence assay (n = 3). (D) HeLa RIPK1/TRADD dKO cells reconstituted with human or mouse RIPK1-GFP were treated with TSZ for 12 h or TCZ for 2 h. Representative confocal fluorescence images show cytosolic RIPK1-GFP puncta and nuclear DAPI staining. Scale bar, 5 μm. The data in all graphs are showed as mean ± SD. Representative immunoblots and immunofluorescence images from three independent experiments are shown. Statistical significance was determined using two-way ANOVA for group comparisons (B, C): ****P < 0.0001.

### RIPK1 kinase inhibitors fail to block kinase-dependent apoptosis in human cells

Given the observed species-specific differences, we evaluated whether RIPK1 kinase inhibitors exhibit differential effects in human and mouse cells. First, we confirmed previous findings that apoptosis induced by TNF/SMAC and TNF/5Z7 is RIPK1 kinase-dependent, whereas TNF/CHX-and TNF/IFNγ-induced apoptosis is RIPK1 kinase-independent. This was demonstrated using kinase-dead knock-in mutations (K45M and D138N) and re-expression of RIPK1 kinase-dead mutant isoforms in RIPK1 knockout cells (Figures 6A-6C and S5A-5G). We next tested five structurally distinct RIPK1 inhibitors (RIPA-56, Nec-1/Nec-1s, GSK2982772, Eclitasertib, and PK68) across a broad panel of human and mouse cells, including both immortalized lines and primary cells. This panel encompassed cell types competent for both necroptosis and kinase-dependent apoptosis, such as human HT29, HaCaT, and U937 cells, human primary macrophages, and mouse iMEF, L929, and BMDM cells. In mouse cells (EMT6, iMEF, L929, and BMDMs), all inhibitors effectively blocked kinase-dependent apoptosis and also suppressed necroptosis in iMEF, L929, and BMDMs (Figures 6D and 6E). A notable exception was PK68, which inhibited kinase-dependent apoptosis in iMEF, EMT6, and L929 cells but not in BMDMs (Figure 6D). The mechanism remains unclear, though it is unlikely to involve BMDM-specific metabolism and inactivation, as PK68 blocked TSZ-induced necroptosis in BMDMs (Figure 6E). In human cells (HT29, HaCaT, human macrophages), inhibitors completely suppressed necroptosis (Figure 6E), but failed to inhibit TNF/SMAC-induced kinase-dependent apoptosis (Figure 6D). The same results were observed in additional human cell lines (HeLa, HCT116, HFF, Hs587T, Jurkat, U937, and MOLT4) (Figure 6D), even after extensive testing across varying TNF or inhibitor concentrations, different treatment durations, and analyzed with various assay types (Figures S5H-S5C1).

**Figure 6.**
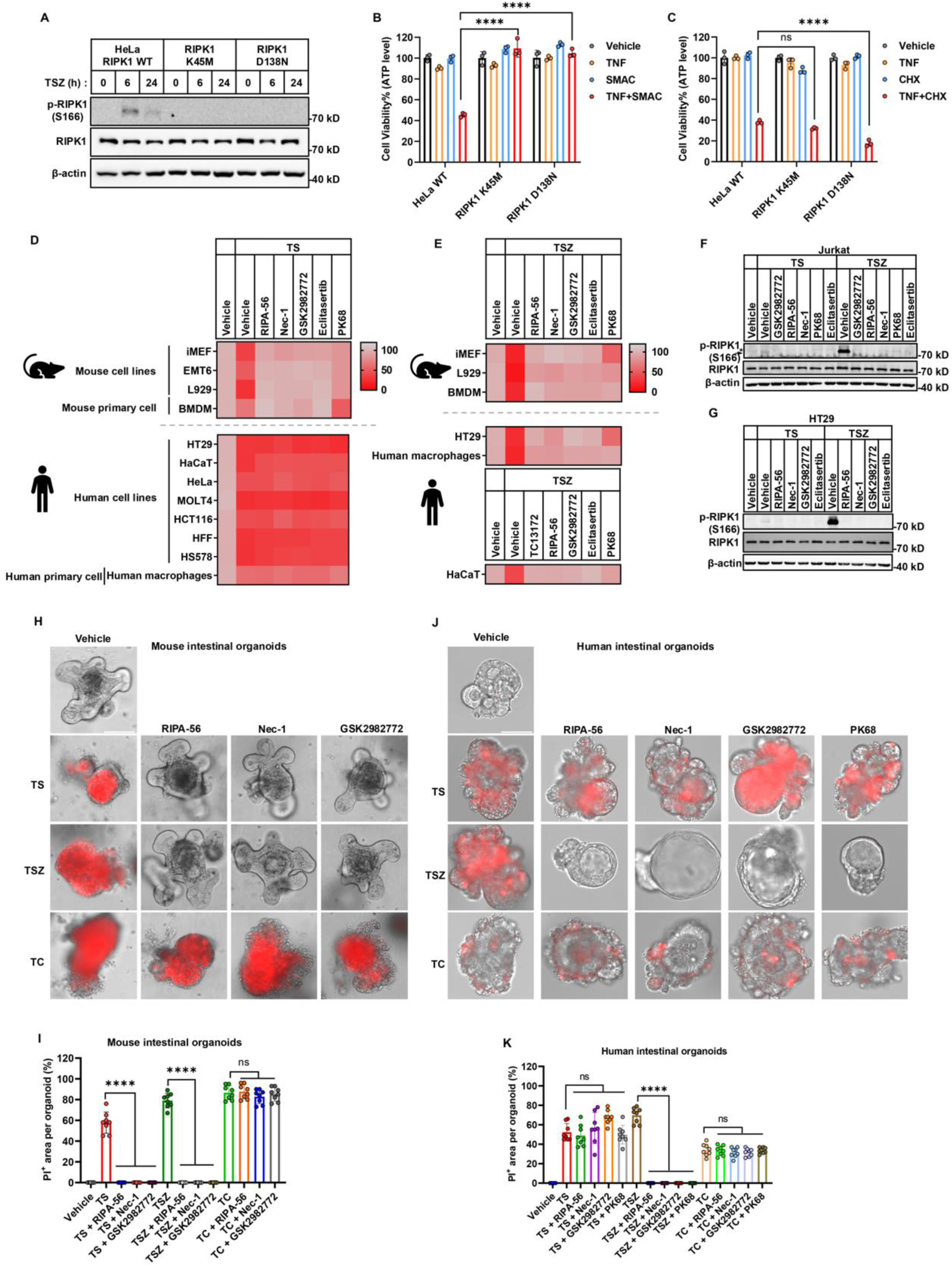
RIPK1 kinase inhibitors block TNF/SMAC-induced apoptosis in mouse cells but not in human cells. (A) HeLa cells (WT, RIPK1-K45M knock-in, and RIPK1-D138N knock-in) were treated with TSZ for 6 or 24 h. Phosphorylated RIPK1 (S166) and total RIPK1 were analyzed by immunoblotting. (B, C) HeLa WT, RIPK1-K45M, and RIPK1-D138N knock-in cells were treated with TNF/SMAC for 48 h (B) or TNF/CHX for 8 h (C), and cell viability was measured by ATP-based luminescence assay (n = 3). Two-way ANOVA: ****P < 0.0001; ns, not significant. Heatmap scaled to the mean of 3 biological replicates. (D, E) Mouse or human cells were treated with TNF/SMAC (D) or TNF/SMAC/z-VAD (E) in the presence or absence of RIPK1 kinase inhibitors (RIPA-56, Nec-1, GSK2982772, Eclitasertib, PK68). Heatmaps showing cell viability following treatment as indicated. Values represent normalized mean cell viability (%) from three biological replicates. Cell viability was measured by ATP-based luminescence assay. (F) Jurkat cells were treated with either TNF/SMAC or TNF/SMAC/z-VAD for 4 h, each in the presence or absence of the indicated RIPK1 kinase inhibitors (RIPA-56, Nec-1, GSK2982772, Eclitasertib or PK68). Phosphorylation of RIPK1 at Ser166 and total RIPK1 levels were analyzed by immunoblotting. Asterisk denotes non-specific bands; arrowhead indicates the phosphorylated band. (G) HT29 cells were treated with either TNF/SMAC or TNF/SMAC/z-VAD for 3 h, in the presence of the indicated RIPK1 inhibitors. Phosphorylation of RIPK1 at Ser166 and total RIPK1 levels were analyzed by immunoblotting. (H, I) Human intestinal organoids were treated with either TNF/SMAC, TNF/SMAC/z-VAD or TNF/CHX, in the presence of indicated RIPK1 inhibitors. Cell death was determined by PI staining (H). Quantification of PI-positive area per organoid corresponding to the fluorescent images (I). n = 8 organoids per condition. Scale bar, 100 μm. (J, K) Mouse intestinal organoids were treated with either TNF/SMAC, TNF/SMAC/z-VAD or TNF/CHX, each in the presence or absence of the indicated RIPK1 inhibitors (RIPA-56, Nec-1, or GSK2982772). Cell death was evaluated by PI staining (J). Quantification of PI-positive area per organoid corresponding to the fluorescent images (K). n = 8 organoids per condition. Scale bar, 100 μm. The data in all graphs are showed as mean ± SD. Representative immunoblots and microscopy images from three independent experiments are shown. Statistical significance was determined using two-way ANOVA (B, C), or one-way ANOVA (I, K) for group comparisons: ****P < 0.0001; ns, not significant.

Thus, while genetic requirements for kinase-dependent apoptosis are similar between species (both RIPK1-dependent, TRADD-independent), pharmacological inhibition of RIPK1 only blocks kinase-dependent apoptosis in mouse cells—not in human cells.

### Species context—not RIPK1 protein—determines inhibitor responsiveness in kinase-dependent apoptosis

To determine whether the insensitivity of human cells to RIPK1 kinase inhibitors in kinase-dependent apoptosis stems from intrinsic resistance of human RIPK1 or from species-specific cellular context, we expressed GFP-tagged or un-tagged human or mouse RIPK1 in RIPK1 KO human cell lines(Figures 7A and 7D). Both forms restored TNF/SMAC-induced apoptosis (Figures 7E, S6B and S6C) and NF-κB activation (Figure S6A). However, RIPK1 kinase inhibitors failed to suppress apoptosis in these cells, regardless of whether human or mouse RIPK1 was expressed (Figures 7B, 7C and 7F).

**Figure 7.**
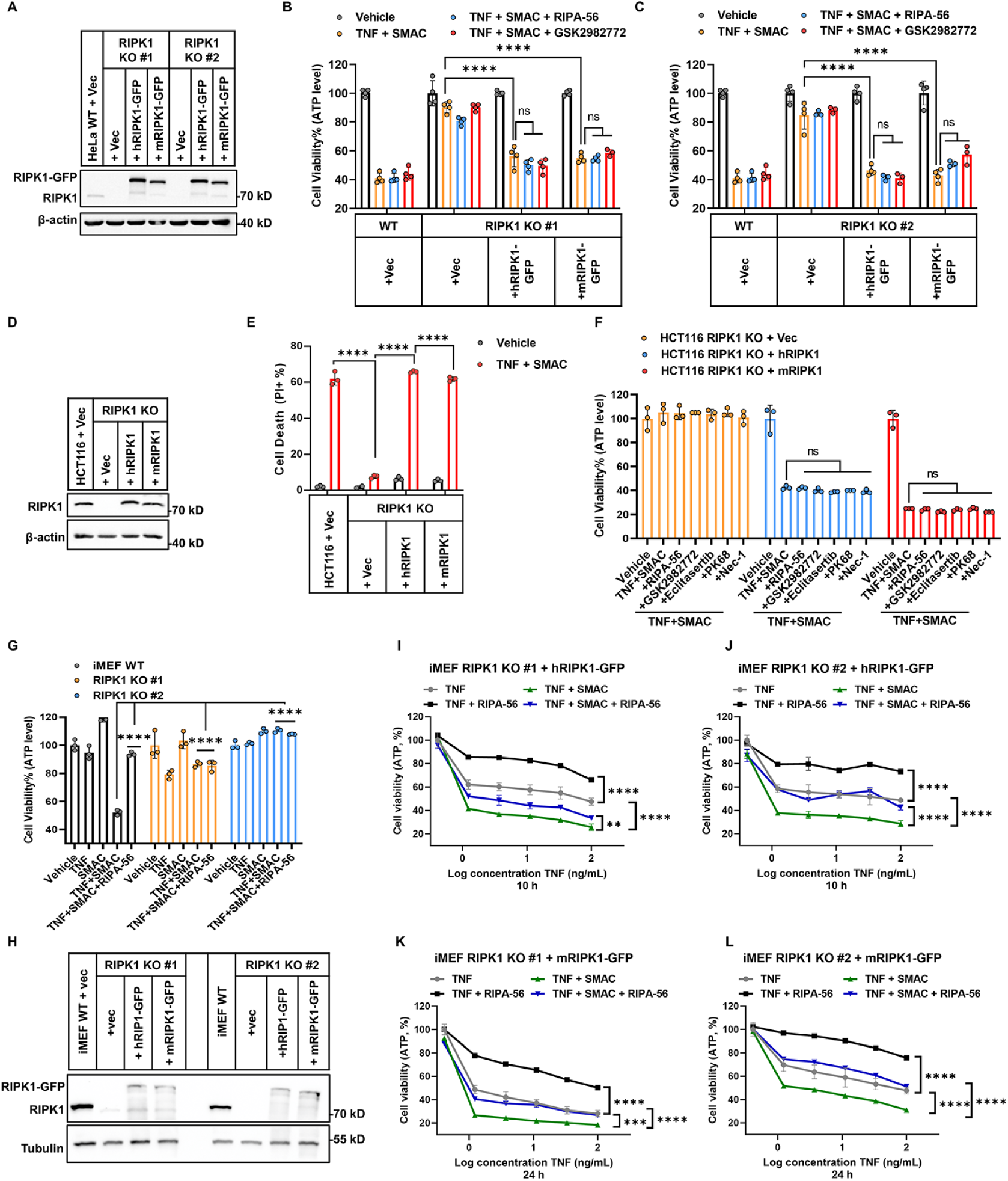
Human and mouse RIPK1 show conserved function in the sensitivity to RIPK1 kinase inhibitors. (A-C) HeLa WT, RIPK1 KO clones #1 and #2, and RIPK1 KO cells reconstituted with human RIPK1-GFP or mouse RIPK1-GFP were analyzed by immunoblotting for RIPK1 expression (A). Cells were treated with TNF/SMAC for 48 h in the presence or absence of RIPK1 kinase inhibitors RIPA-56 or GSK2982772. Cell viability was measured by ATP-based luminescence assay in clone #1 (B) and clone #2 (C) (n = 3). (D-F) HCT116 WT, RIPK1 KO, and RIPK1 KO cells reconstituted with human RIPK1 or mouse RIPK1 were analyzed by immunoblotting for RIPK1 expression (D). Cells were treated with TNF/SMAC (E), or with TNF/SMAC in the presence or absence of RIPK1 kinase inhibitors for 24 h (F). Cell death was quantified by propidium iodide (PI) staining followed by flow cytometry (E), cell viability was assessed by ATP-based luminescence assay (F) (n = 3). (G) iMEF WT, RIPK1 KO clones #1 and #2 were treated with TNF/SMAC in the presence or absence of RIPA-56 for 24 h. Cell viability was assessed by ATP-based luminescence assay (n = 3). (H) RIPK1 expression was analyzed by immunoblotting in iMEF WT, RIPK1 KO clones #1 and #2 reconstituted with human RIPK1-GFP or mouse RIPK1-GFP. (I-L) iMEF RIPK1 KO clones #1 and #2 reconstituted with human RIPK1-GFP (I, J) or mouse RIPK1-GFP (K, L) were treated with TNF or TNF/SMAC in the presence or absence of RIPA-56. Cell viability was assessed by ATP-based luminescence assay (n = 3). The data in all graphs are showed as mean ± SD. Representative immunoblots from three independent experiments are shown. Statistical significance was determined by two-way ANOVA (B, C, E, G), or one-way ANOVA (F, I-L) for group comparisons: ****P < 0.0001, ***P < 0.001 (K, P=0.0009), **P < 0.01 (I, P=0.0033); ns, not significant.

To further dissect the role of cellular context, we introduced human or mouse RIPK1 into murine iMEF RIPK1 KO cells (Figures 7G and 7H). Paradoxically, reconstitution with either RIPK1 variant in two independent clones sensitized cells to TNF-induced apoptosis in a dose-dependent manner (Figures 7I-7L). And co-treatment with a sub-lethal dose of SMAC further enhanced cell death (Figures 7I-7L). Notably, TNF-or TNF/SMAC-induced apoptosis could be partially rescued by RIPK1 inhibitor in both human-and mouse-RIPK1–reconstituted iMEF cells at comparable levels, indicating functional equivalence of the two RIPK1 orthologs (Figures 7I-7L). These findings collectively demonstrate that the differential inhibitor sensitivity is dictated by species-specific cellular context rather than by intrinsic properties of the RIPK1 protein.

Because TRADD can support kinase-independent apoptosis, we tested whether RIPK1 inhibitors might switch kinase-dependent apoptosis to a TRADD-mediated form in human cells. However, in HeLa TRADD KO, MCF-7 TRADD KO, and HaCaT TRADD KO cells, RIPK1 inhibitors failed to block TNF/SMAC-induced apoptosis, similar to the corresponding WT cells (Figures S6D-S6F). Likewise, in RIPK1/TRADD dKO HeLa cells reconstituted with either human or murine RIPK1, inhibitors still did not block TNF/SMAC-induced apoptosis (Figure S6H). In HaCaT cells, where TSZ triggered necroptosis, RIPK1 inhibitors blocked necroptosis, serving as a control for effective inhibitor activity (Figure S6G). These results indicate that TNF/SMAC-induced apoptosis in the presence of RIPK1 inhibitors does not arise from a switch to TRADD-dependent death.

Since RIPK1 inhibitors block necroptosis but spare apoptosis in human cells, we next tested whether this was due to incomplete inhibition of RIPK1 kinase activity. We measured autophosphorylation of RIPK1 S166, a marker of kinase activity, in Jurkat and HT29 cells undergoing TNF/SMAC-induced apoptosis or TSZ-induced necroptosis. TNF/SMAC induced weaker S166 phosphorylation than TSZ, but in both contexts phosphorylation was efficiently blocked by RIPK1 inhibitors (Figures 6F and 6G). Thus, the inability of inhibitors to prevent apoptosis in human cells is not due to a failure to block RIPK1 S166 autophosphorylation.

### Implications of inhibitor responsiveness to therapeutic translation

Our finding that RIPK1 inhibitors block kinase-dependent apoptosis in mouse but not in human cells parallels the clinical outcomes of these compounds. RIPK1 inhibitors show strong efficacy in multiple mouse inflammatory models—including ulcerative colitis, rheumatoid arthritis, psoriasis, Alzheimer’s disease, multiple sclerosis, and amyotrophic lateral sclerosis^58–69^—yet corresponding human clinical trials have failed (trials NCT05630547 and NCT05237284).^39–42,70,71^ To date, ∼20 RIPK1 inhibitors have entered clinical development (https://www.pharmacodia.com/), but none has achieved regulatory approval, and no publicly reported trials have met primary efficacy endpoints.

To strengthen human relevance beyond cultured cells, we included *ex vivo* intestinal organoids from both species. RIPK1 inhibitors blocked TSZ-induced necroptosis in both systems, confirming efficient inhibitor activity (Figures 6H-6K). Consistent with cultured cells, RIPK1 inhibitors (RIPA-56, Nec-1, GSK2982772) blocked TNF/SMAC-induced apoptosis in mouse organoids but failed in human organoids (Figures 6H-6K). Collectively, results from primary macrophages and intestinal organoids, along with findings from immortalized cell lines, suggest that the lack of efficacy of RIPK1 inhibitors in human kinase-dependent apoptosis may underlie their repeated clinical failures.

## Discussion

Research on the TNF pathway has relied heavily on both *in vitro* mouse cells and *in vivo* mouse models. In many recent reviews, signaling pathways have been presented by merging findings from human and mouse cells indiscriminately, shaping the field’s narrative, simplifying the biology for non-specialists, and guiding translation from mouse studies into clinical trials.^9,26,55,56^ All of this rests on the assumption that TNF signaling is conserved between humans and mice. However, repeated clinical failures of RIPK1 inhibitors that were highly effective in mouse models challenge this assumption. ^39–42,70,71^ Against this background, our study systematically examined the roles of RIPK1 and TRADD in TNF signaling, including NF-κB and MAPK activation, kinase-dependent and-independent apoptosis, necroptosis, and sensitivity to RIPK1 inhibitors (Figure S7). We uncovered fundamental species-specific differences in both apoptosis modalities.

Specifically, RIPK1 inhibitors blocked kinase-dependent apoptosis in mouse but not in human cells, despite efficiently inhibiting RIPK1 S166 autophosphorylation. Genetic approaches (K45M and D138N mutations) confirmed the essential role of RIPK1 kinase activity in apoptosis. These results suggest that inhibitor-bound RIPK1 may retain partial kinase activity during TNF/SMAC stimulation in human cells. Importantly, apoptosis in the presence of RIPK1 inhibitors did not switch to TRADD-dependent death. Cross-species re-expression of human and mouse RIPK1 further demonstrated that the discrepancy arises from cell context rather than RIPK1 itself. Future work on differentially bound proteins, novel substrates, or alternative autophosphorylation sites of RIPK1 will help clarify the mechanism.

For kinase-independent apoptosis, RIPK1 and TRADD act redundantly in human cells, with apoptosis blocked only when both are deleted. In contrast, in mouse cells RIPK1 KO facilitated apoptosis, whereas TRADD KO completely blocked it, regardless of RIPK1 status (Figure S7C). Expression of mouse RIPK1 in human RIPK1/TRADD dKO cells restored apoptosis to the same extent as human RIPK1, again pointing to cell context as the determinant. Similarly, comparative analyses of differentially bound proteins and post-translational modifications of RIPK1 between human and mouse cells will help narrow down potential factors and provide insights into the nature of this cell context. Intriguingly, the same species-specific factors that support RIPK1’s role in kinase-independent apoptosis may also underlie the insensitivity of human cells to RIPK1 kinase inhibitors.

Our discovery that RIPK1 KO in mouse cells facilitates kinase–independent, and TRADD-dependent apoptosis is consistent with phenotypes observed in genetic RIPK1 KO mice.^15,72,73^ During embryogenesis, RIPK1 deficiency triggers apoptosis and necroptosis in multiple tissues, leading to lethality late in gestation or shortly after birth.^72–75^ Specifically, RIPK1 KO promotes apoptosis in mouse tissues through TRADD-mediated formation of complex II with FADD and caspase-8.^15^ When TRADD is also knocked out, RIPK1 deficiency–mediated activation of caspase-8 and caspase-3 is largely suppressed. Therefore, the essential role of TRADD (somehow inhibited by RIPK1) in kinase-independent apoptosis, as observed in our *in vitro* study, is consistent with prior *in vivo* findings in mice.^15^

Cell type–specific factors inevitably influence biological processes. Indeed, when testing RIPK1 inhibitors in additional cell lines, some minor inconsistencies emerged. For example, in Jurkat cells all RIPK1 inhibitors provided mild but significant protection against kinase-dependent apoptosis (Figure S5Q). In HeLa cells, Nec-1 protection was generally more pronounced than in other lines, especially when measured by LDH release (Figure S5H). The choice of small molecules—particularly Nec-1/Nec-1s—also had some impact, likely due to their known off-target effects,^76^ which can lead to misinterpretation. To minimize such confounding factors, we used five structurally distinct, clinically relevant RIPK1 inhibitors (Nec-1/Nec-1s, RIPA-56, GSK2982772, Eclitasertib, PK68), yielding consistent results that strengthen our conclusions.

Our findings appear to conflict with a published study (and potentially other studies) reporting that Nec-1s reduced LDH release upon TNF/BV6 stimulation in HeLa cells.^77^ We attribute this discrepancy to a combination of factors, including off-target effects of Nec-1s, choice of assay (LDH release), cell type (HeLa), and death trigger (TNF/BV6 vs. TNF/SMAC) as above mentioned (Figures S5H-S5J). Importantly, the same study showed that Nec-1s only partially protected TNF/BV6-induced apoptosis in MIB2 KO HeLa cells and failed to protect MIB2 KO HCT116 cells, supporting our conclusion. ^77^

Because we systematically re-examined TNF downstream signaling, some overlap with previous reports was unavoidable. However, the novelty of our study lies not only in generating new conclusions but also in the unexpected, systematic identification of multiple species-specific differences between human and mouse cells. This is critically important in a field that relies heavily on mouse models for preclinical studies and clinical translation. The species-specific discrepancies we uncovered highlight the need to revise current assumptions, provide insight into the clinical failure of RIPK1 inhibitors, and suggest new strategies—either developing inhibitors effective against both kinase-dependent apoptosis and necroptosis in human cells, or targeting alternative targets to simultaneously block kinase-dependent and-independent death. Our findings also argue for the use of humanized mouse models in studying inflammatory diseases.

## Materials and methods

### Cell culture

All mammalian cells were maintained at 37 °C in a humidified incubator with 5% CO₂. HeLa, HCT116, MCF-7, HaCAT, HFF (human foreskin fibroblast), iMEF, BMDM, L929, Hs578T, and 293T cells were cultured in Dulbecco’s Modified Eagle Medium (DMEM; Gibco, C11995500BT). EMT6, CT26, Jurkat, U937, human macrophages, and MOLT4 cells were cultured in RPMI-1640 medium (Gibco, C11875500BT). HT29 cell was cultured in McCoy’s 5A (Procell; PM150710P). HeLa and HCT116 cell lines were obtained from the American Type Culture Collection (ATCC). Other cells were obtained from the Cell Resource Center, Peking Union Medical College (PCRC). All cell lines were routinely tested and confirmed to be free of mycoplasma contamination.

### Treatment of cells with cytokines and compounds

Different cytokine and compound treatments were applied to various cell lines. HeLa, HaCAT, HFF, human macrophages, MOLT4, Jurkat, U937, Hs578T, HT29, and MCF-7 cells were treated with 50 ng/mL TNF (Cat# 10602-HNAE, Sino Biological) in combination with either 500 nM Smac mimetic (CAS No.: 411230-24-5; synthesized in-house), 25 μg/mL cycloheximide (CHX, Cat# 239763-M, Sigma-Aldrich), 100 ng/mL IFNγ (Cat# 11725-HNAS, Sino Biological), 1-10μM BV6 (Cat# BD629814, Bidepharm), or 250 nM 5Z7 (Cat# O9890, Sigma-Aldrich). HCT116 cells were treated with 2 ng/mL TNF plus 500 nM Smac, or 50 ng/mL TNF plus 25 μg/mL CHX. EMT6 and iMEF cells were treated with 50 ng/mL TNF combined with 500 nM Smac, 25 μg/mL CHX, 100 ng/mL IFNγ, or 250 nM 5Z7. Intrinsic apoptosis was induced by treating cells with 5 μM ABT-737 (Cat# 197333, Sigma-Aldrich) and 2 μM S63845 (Cat# HY-100741, MedChemExpress). Necroptosis was induced with 50 ng/mL TNF, 500 nM Smac mimetic, and 20 μM z-VAD-FMK (Cat# GC12861, GlpBio). RIPK1 kinase inhibitors used in this study included compound RIPA-56 (synthesized in-house),^78^ Nec-1 (Cat# HY-15760, MedChemExpress), GSK2982772 (Cat# HY-101760, MedChemExpress), Eclitasertib (Cat# HY-114371, MedChemExpress), Nec-1s (Cat# C275378, Aladdin), and PK68 (Cat# HY-128348, MedChemExpress). For MLKL inhibition, compound TC13172 (synthesized in-house)^79^ was used.

### Cell viability assay

Cell viability was assessed by measuring intracellular ATP levels using the CellTiter-Glo Luminescent Cell Viability Assay (Cat# G7573; Promega), following the manufacturer’s protocol. Luminescence was detected with a PerkinElmer EnSpire microplate reader. Data were processed using GraphPad Prism (GraphPad Software, San Diego, CA, USA), and dose–response curves were fitted using a non-linear regression model with a sigmoidal (variable slope) function.

### Cell death assay by PI staining

A total of 1 × 10⁵ cells per well were seeded in 24-well plates one day prior to treatment. Cells were then subjected to the indicated treatments for the designated durations. For flow cytometry analysis, cells were harvested by trypsinization and resuspended in PBS containing 1 μM propidium iodide (PI; Cat# 537059, Sigma-Aldrich). After a 10-minute incubation at room temperature, samples were analyzed using a flow cytometer, and data were processed with FlowJo software. For live-cell imaging, cells were washed three times with PBS and incubated in fresh culture medium containing 1 μM PI for 10 minutes. Images were acquired using a Thermo EVOS M7000 imaging system.

### LDH release

Plasma membrane rupture (PMR) was assessed by measuring lactate dehydrogenase (LDH) released into the culture supernatant using the CytoTox 96 Non-Radioactive Cytotoxicity Assay (Promega; G1780), following the manufacturer’s instructions. Absorbance at 490 nm was measured using a microplate reader (PerkinElmer EnSpire), and the data were analyzed using GraphPad Prism (GraphPad Software, Inc., San Diego, CA, USA).

### Western blotting

Cells were lysed either in RIPA buffer (50 mM Tris-HCl, pH 7.4; 150 mM NaCl; 1% Triton X-100; 1% sodium deoxycholate; 0.1% SDS) supplemented with cOmplete™ EDTA-free Protease Inhibitor Cocktail (Cat# 05892791001; Roche), or directly solubilized in 1× SDS sample buffer. For RIPA lysis, samples were incubated on ice for 30 minutes and centrifuged at 20,000 × g for 10 minutes at 4 °C. Supernatants were then mixed with 4× SDS loading buffer and subjected to SDS–PAGE. The following antibodies were used for immunoblotting: anti-HSP90 (13171-1-AP; Proteintech), anti-caspase-8 (4790; Cell Signaling Technology), anti-β-actin (4970; Cell Signaling Technology), anti-GAPDH (HRP-60004; Proteintech), anti-SHARPIN (14626-1-AP; Proteintech), anti-p-RIPK1 (S166) (31122; Cell Signaling Technology), anti-RIPK1 (3493; Cell Signaling Technology), anti-FADD (14906-1-AP; Proteintech), anti-TNFR1 (21574-1-AP; Proteintech), anti-TRADD (15468-1-AP; Proteintech), anti-IκBα (4814; Cell Signaling Technology), anti-p-NF-κB p65 (3033; Cell Signaling Technology), anti-NF-κB p65 (8242; Cell Signaling Technology), anti-p-p38 MAPK (4511; Cell Signaling Technology), anti-p38 MAPK (14064-1-AP; Proteintech), anti-tubulin (10068-1-AP; Proteintech), anti-FLAG (20543-1-AP; Proteintech), anti-Myc (16286-1-AP; Proteintech), HRP-conjugated anti-rabbit IgG (A0545; Sigma-Aldrich), and HRP-conjugated anti-mouse IgG (A9044; Sigma-Aldrich).

### Stable Cell Line Generation

Lentiviral particles were produced by co-transfecting 293T cells with the expression plasmids (encoding human RIPK1, mouse RIPK1, human TRADD, or mouse TRADD), packaging plasmid psPAX2, and envelope plasmid pMD2.G, using polyethylenimine (PEI; 24765, Polysciences). After 48 hours, the viral supernatant was harvested and filtered, then used to transduce target cells in the presence of 5 μg/mL Polybrene (C0351, Beyotime). Twenty-four hours post-transduction, cells were selected with either 10 μg/mL puromycin (ST551, Beyotime) or 10 μg/mL blasticidin (BLL-45-06, InvivoGene) for 7 consecutive days to establish stable cell lines.

### CRISPR–Cas9-mediated gene knockouts

Gene knockout cell lines were generated using CRISPR-Cas9 technology. HeLa, HCT116, MCF-7, EMT6, or iMEF cells were transfected with sgRNA-expressing pX458 plasmids using Lipofectamine 3000 (Thermo Fisher Scientific) according to the manufacturer’s protocol. GFP-positive cells were sorted by flow cytometry using a BD FACSAria Fusion cell sorter. Single GFP-positive cells were seeded into 96-well plates for clonal expansion. Once confluent, clones were transferred to 24-well plates, and successful knockouts were confirmed by immunoblotting. A complete list of all primers used can be found in Supplementary Table 1.

### Knockin mutation of RIPK1-K45M, D138N in HeLa cell

To generate RIPK1-K45M and RIPK1-D138N knock-in mutations in HeLa cells, single-stranded oligodeoxynucleotides (ssODNs) containing the desired mutations flanked by 40-nucleotide homology arms were designed. The ssODN sequences were as follows: K45M (ssODN):5’-GGAAGGTGTCTCTGTGTTTCCACAGAACCCAGGGACTCATGATCATGATGACAGTGTACAAGGGCCCCAACTGCATTGAGTGAGTAGGGAGCAGGCGTGG-3’,D138N (ssODN) 5’-TCATTGAAGGAATGTGCTACTTACATGGAAAAGGCGTGATACACAAA AAC CTGAAGCCTGAAAATATCCTTGTTGATAATGACTTCCACATTAAGGTAAA-3’. The mutated nucleotides are underlined.

To introduce the mutations, sgRNAs targeting the respective loci (K45 sgRNA: 5’-CATGAAAACAGTGTACAAG-3’; D138 sgRNA: 5’ - GGAAAAGGCGTGATACACA-3′) were cloned into the pX458 vector. HeLa cells were co-transfected with 10 nM pX458-sgRNA and 10 nM ssODN using Lipofectamine 3000 (Thermo Fisher Scientific). To enhance homology-directed repair (HDR) efficiency, 20 µM NU7026 (N1537; Sigma-Aldrich) was included during transfection to inhibit non-homologous end joining (NHEJ). Single GFP-positive cells were sorted by flow cytometry, expanded, and screened by Sanger sequencing to confirm the desired mutations.

### Complex I immunoprecipitation

HeLa cells or MEFs were pre-treated as described in the figure legends and subsequently stimulated, or left unstimulated, with 200 ng/mL FLAG-tagged human TNF (FLAG-hTNF) or mouse TNF (FLAG-mTNF) for the indicated durations. Cells were washed twice with ice-cold PBS and lysed in NP-40 lysis buffer (10% glycerol, 0.5% NP-40, 150 mM NaCl, 10 mM Tris-HCl, pH 8.0) supplemented with EDTA-free protease inhibitor cocktail (4693132001; Roche). Lysates were cleared by centrifugation at 20,000 × g for 10 minutes at 4°C, and the supernatants were incubated overnight with ANTI-FLAG M2 affinity gel (A2220; Sigma-Aldrich). On the following day, the beads were washed four times with NP-40 lysis buffer. Immunoprecipitates or whole-cell lysates were then resuspended in 1× SDS sample buffer and analyzed by SDS-PAGE followed by immunoblotting.

### Complex II immunoprecipitation

FADD-GFP–expressing iMEFs were pre-treated as indicated in the figure legends for the specified durations. Cells were washed twice with ice-cold PBS and lysed in NP-40 lysis buffer (10% glycerol, 0.5% NP-40, 150 mM NaCl, 10 mM Tris-HCl, pH 8.0) supplemented with EDTA-free protease inhibitor cocktail (4693132001; Roche). Cell lysates were cleared by centrifugation at 20,000 × g for 10 minutes at 4°C. The resulting supernatants were incubated overnight at 4°C with anti-GFP magnetic beads. The next day, beads were washed four times with NP-40 lysis buffer. Immunoprecipitates or input lysates were then resuspended in 1× SDS sample buffer and subjected to SDS-PAGE followed by immunoblot analysis.

### Immunofluorescence

HeLa and MEF cells stably expressing FADD-GFP were pre-treated as described in the figure legends for the indicated time. Cells were then fixed with 4% paraformaldehyde for 15 minutes at room temperature, followed by three washes with PBS. Nuclei were stained with Hoechst (C0031; Solarbio) diluted in PBS for 10 minutes, and samples were again washed three times with PBS. Coverslips were stored at 4°C prior to imaging. Fluorescence images were acquired using a Zeiss LSM 980 confocal microscope with a 63× oil immersion objective and analyzed using Zeiss Zen software. Manders’ overlap coefficient was calculated using ImageJ. All images shown are representative of at least three independent experiments.

### Isolation and differentiation of human primary macrophages

Human monocyte-derived macrophages were generated from PBMCs (YaYuBio; CWPB025M). PBMCs were cultured in RPMI-1640 medium supplemented with 10% heat inactivated FBS (2.5×10⁷ cells per 15 cm plate). To differentiate the PBMCs into macrophages, macrophage colony-stimulating factor (M-CSF, 20 ng/mL; Sino Biological, 11792-HNAH) was added to the medium for 5 days. On day 6, the differentiated macrophages were washed with cold PBS, scraped from the plates, and centrifuged at 500 g for 5 minutes. The cells were then resuspended in 24-well plates at a density of 5×10⁵ cells per well. After 12 h, the cells were treated with TNF/SMAC, or TNF/SMAC/z-VAD.

### Isolation and Culture of Bone Marrow-Derived Macrophages (BMDMs)

Bone marrow-derived macrophages (BMDMs) were isolated from the femurs and tibias of 6-to 8-week-old C57BL/6J mice. All animal procedures were approved by the Institutional Animal Care and Use Committee of the Institute of Genetics and Developmental Biology, Chinese Academy of Sciences. Bone marrow cells were flushed with sterile ice-cold PBS, passed through a 70-μm strainer, and subjected to red blood cell lysis (Cat# C3702, Beyotime). Cells were plated in non-tissue-culture-treated Petri dishes and differentiated for 7 days at 37 °C, 5% CO₂ in complete RPMI-1640 containing 30% (v/v) L929 cell conditioned medium, 10% heat-inactivated FBS, and 1% penicillin/streptomycin. On day 3, half of the medium was gently replaced with fresh complete medium containing 30% L929 cell conditioned medium; an optional half-medium refresh was performed on day 6. On day 7, adherent BMDMs were harvested using cold PBS and gentle scraping for subsequent experiments. The cells were then resuspended in 24-well plates at a density of 5×10⁵ cells per well. After 12 h, the cells were treated with TNF/SMAC, or TNF/SMAC/z-VAD.

### Isolation and Culture of Intestinal Organoids

Mouse small intestinal crypts were isolated from 6-8-week-old C57BL/6J mice. The entire small intestinal was excised, opened longitudinally, and washed thoroughly in cold PBS. The tissue was then fragmented and incubated in PBS containing 2 mM EDTA for 30 minutes on ice with gentle agitation. Crypts were released by vigorous shaking in cold PBS. The supernatant containing isolated crypts was filtered through a 70-μm cell strainer and centrifuged. The crypt pellet was resuspended in Matrigel and plated as droplets in a pre-warmed cell culture plate. After polymerization at 37°C for 20 minutes, the Matrigel domes were overlaid with complete IntestiCult™ Organoid Growth Medium (STEMCELL Technologies) or a custom medium containing EGF, Noggin, and R-spondin-1. The culture medium was refreshed every 2-3 days. Organoids were routinely passaged every 5-7 days by mechanical dissociation and re-embedded in Matrigel. the intestinal organoids were treated with TNF/SMAC, TNF/SMAC/z-VAD, or TNF/CHX.

Human ileum organoids were purchased from AIMINGMED (Hangzhou) and The Third People’s Hospital of Chengdu, the culture method is the same as that for mouse intestinal organoids.

## Ethics declarations

All animal experiments involving mouse tissues were conducted according to protocols approved by the Ethics Committee of the Institute of Genetics and Developmental Biology, Chinese Academy of Sciences, Beijing, China (permit number: AP2025044). Human peripheral blood mononuclear cells (PBMCs) were used in this study and were purchased from Yayu Biotechnology Co., Ltd, Shanghai, China. Human ileum organoids were purchased from AIMINGMED (Hangzhou, China). No personal data or identifiable human information was used, and the cell lines were not derived from individuals involved in clinical trials or research requiring specific consent. The use of human cells complies with all applicable ethical guidelines and regulations and was approved by the Ethics Committee of the Institute of Genetics and Developmental Biology, Chinese Academy of Sciences (permit number: IGDB-2025-IRB-009).

## Statistical analysis

Statistical analysis is performed using GraphPad Prism 8.0 software. The results are presented as mean ± SD from three independent experiments. All of the statistical details can be found in the figure legends.

## Acknowledgements

We thank Dr. Lilin Du at the National Institute of Biological Sciences, Beijing, for providing the beads used in GFP pulldown. This work was supported by the National Key R&D Program of China (2022YFA1304500 to Y.A.); the National Natural Science Foundation of China (22477095, 22307139 to B.Y.; 32321004, 32422023 to Y.A.); and the Beijing Natural Science Foundation Changping Frontier Project (L244054 to Y.A.).

## Author contributions

B.Y. and Y.A. conceptualized and supervised the project. Z.D., B.D., and J.W. conducted the majority of the experiments. K.Y. assisted with validation of gene knockouts and western blot analysis. H.L. contributed to the purification of cytokines required for cell-based assays. The manuscript was written by B.Y., Y.A., and Z.D., with input from all authors.

## Data availability

All the data supporting the findings of this study are available upon request from the corresponding authors.

---

**Figure S1.**
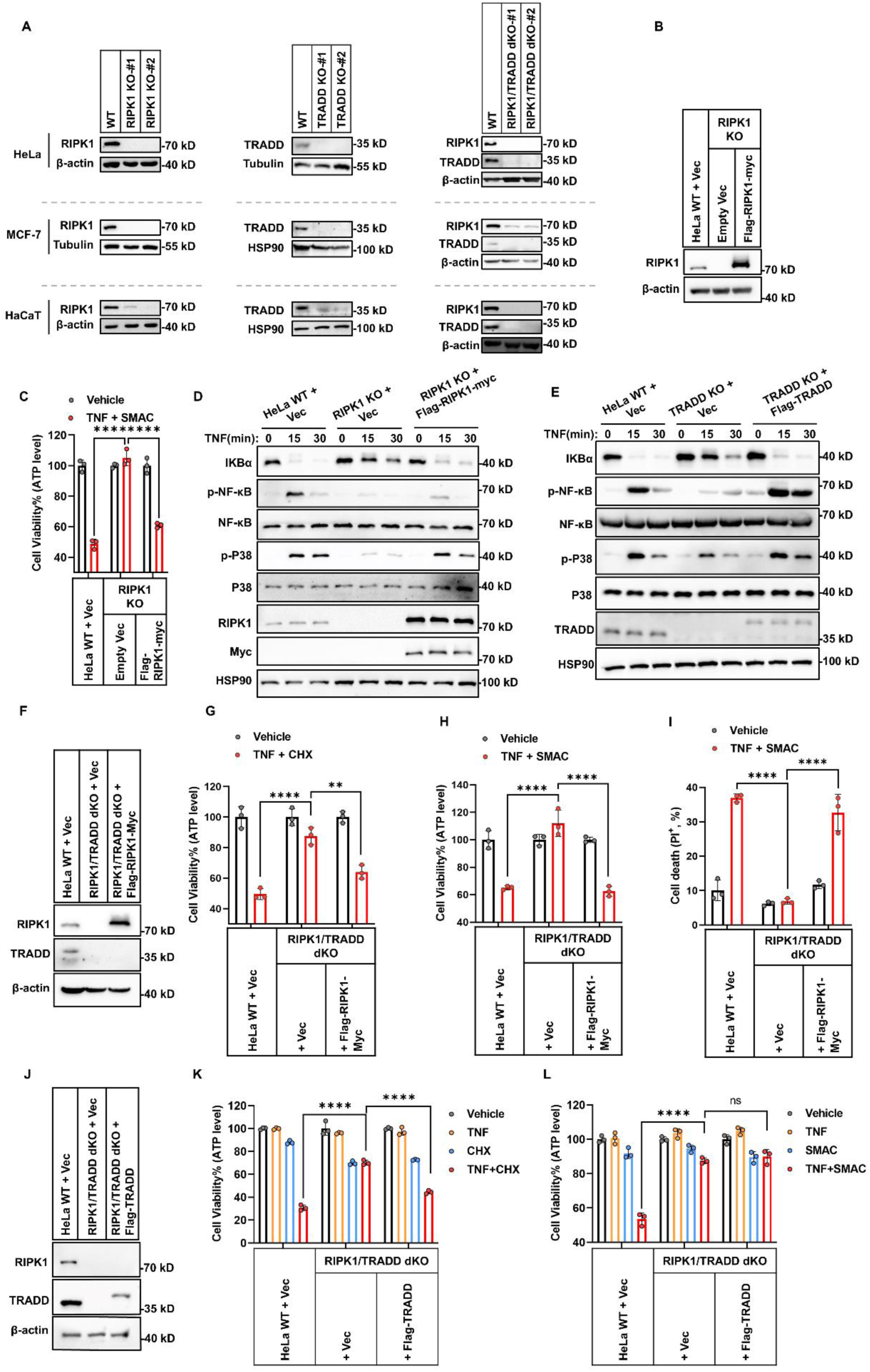
Functional reconstitution of RIPK1 or TRADD restores TNF-induced cell death and NF-κB activation in HeLa knockout cells. (A) RIPK1 KO, TRADD KO, RIPK1 and TRADD dKO in HeLa, MCF-7 or HaCaT cells (clones #1 and #2 from two different gRNAs) was confirmed by immunoblotting. (B-D) RIPK1 KO HeLa cells were reconstituted with RIPK1. RIPK1 expression in WT, KO, and reconstituted cells were confirmed by immunoblotting (B). Cells were treated with TNF/SMAC for 48 h, and cell viability was measured (C) (n = 3). For pathway analysis, cells were stimulated with TNF for 0, 15, or 30 min and assessed for NF-κB and MAPK signaling activation by immunoblotting (D). (E) TRADD KO HeLa cells reconstituted with wild-type TRADD were treated with TNF for the indicated time points. Expression of NF-κB and MAPK pathway components was confirmed by immunoblotting. (F-I) HeLa WT, RIPK1/TRADD dKO, and dKO cells stably reconstituted with RIPK1 were analyzed by immunoblotting (F). Cells were treated with TNF/CHX for 8 h (G) or TNF/SMAC for 48 h (H, I). Cell viability was measured by ATP-based luminescence assay (G, H), and cell death was measured by PI staining (I) (n = 3). (J-L) HeLa WT, RIPK1/TRADD dKO, and dKO cells stably reconstituted with TRADD were analyzed by immunoblotting (J). Cells were treated with TNF/CHX for 8 h (K) or TNF/SMAC for 48 h (L), and cell viability was measured by ATP-based luminescence assay (n = 3). The data in all graphs are showed as mean ± SD. Representative immunoblots from three independent experiments are shown. Statistical significance was determined using two-way ANOVA for group comparisons (C, G-I, K, L): ****P < 0.0001, **P < 0.01 (G, P=0.0012), ns: not significant.

**Figure S2.**
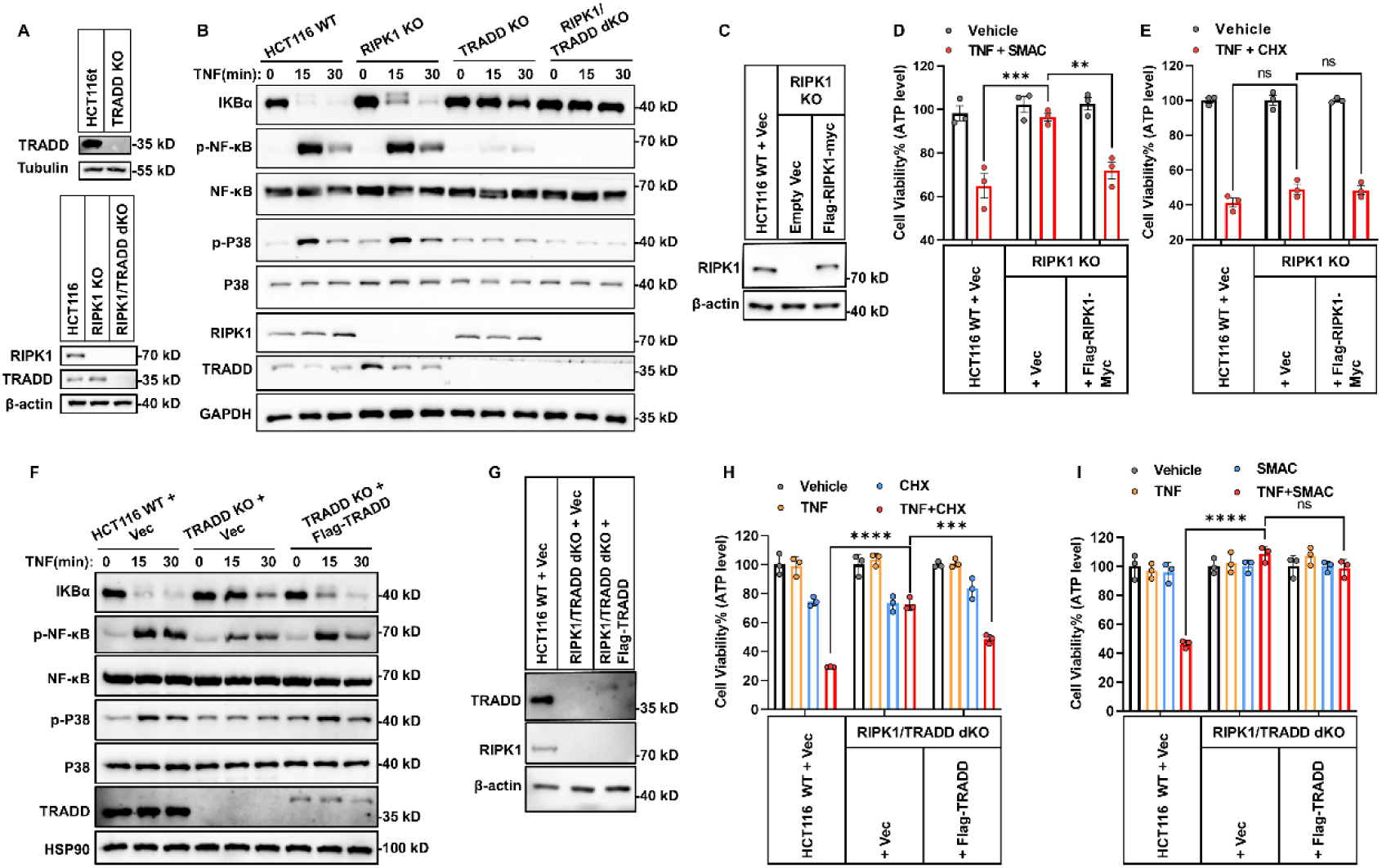
RIPK1 and TRADD are functionally redundant in TNF signaling in HCT116 cells. (A) RIPK1 KO, TRADD KO, RIPK1/TRADD dKO in HCT116 cells was confirmed by immunoblotting. (B) WT, RIPK1 KO, TRADD KO, and RIPK1/TRADD dKO cells were treated with TNF for 0, 15, or 30 minutes. NF-κB and MAPK pathway activation was evaluated by immunoblotting for key signaling proteins (n = 3). (C) RIPK1 expression was confirmed by immunoblotting in HCT116 WT, RIPK1 KO, and RIPK1 KO cells reconstituted with RIPK1. (D, E) HCT116 WT, RIPK1 KO, and RIPK1-reconstituted cells were treated with TNF/SMAC for 24 h (D) or TNF/CHX for 8 h (E), and cell viability was measured by ATP-based luminescence assay (n = 3). (F) HCT116 WT, TRADD KO, and TRADD-reconstituted cells were treated with TNF for 0, 15, or 30 min, and NF-κB/MAPK pathway activation was analyzed by immunoblotting. (G) RIPK1 and TRADD expression was confirmed by immunoblotting in HCT116 WT, RIPK1/TRADD dKO, and TRADD-reconstituted dKO cells. (H, I) Cells were treated with TNF/CHX for 8 h (H) or TNF/SMAC for 24 h (I), and cell viability was assessed using ATP-based luminescence assay (n = 3). The data in all graphs are showed as mean ± SD. Representative immunoblots from three independent experiments are shown. Statistical significance was determined using two-way ANOVA for group comparisons (D, E, H, I): ****P < 0.0001; ***P < 0.001 (D, P=0.0007 and H, P=0.0002); **P < 0.01 (D, P=0.0059); ns, not significant.

**Figure S3.**
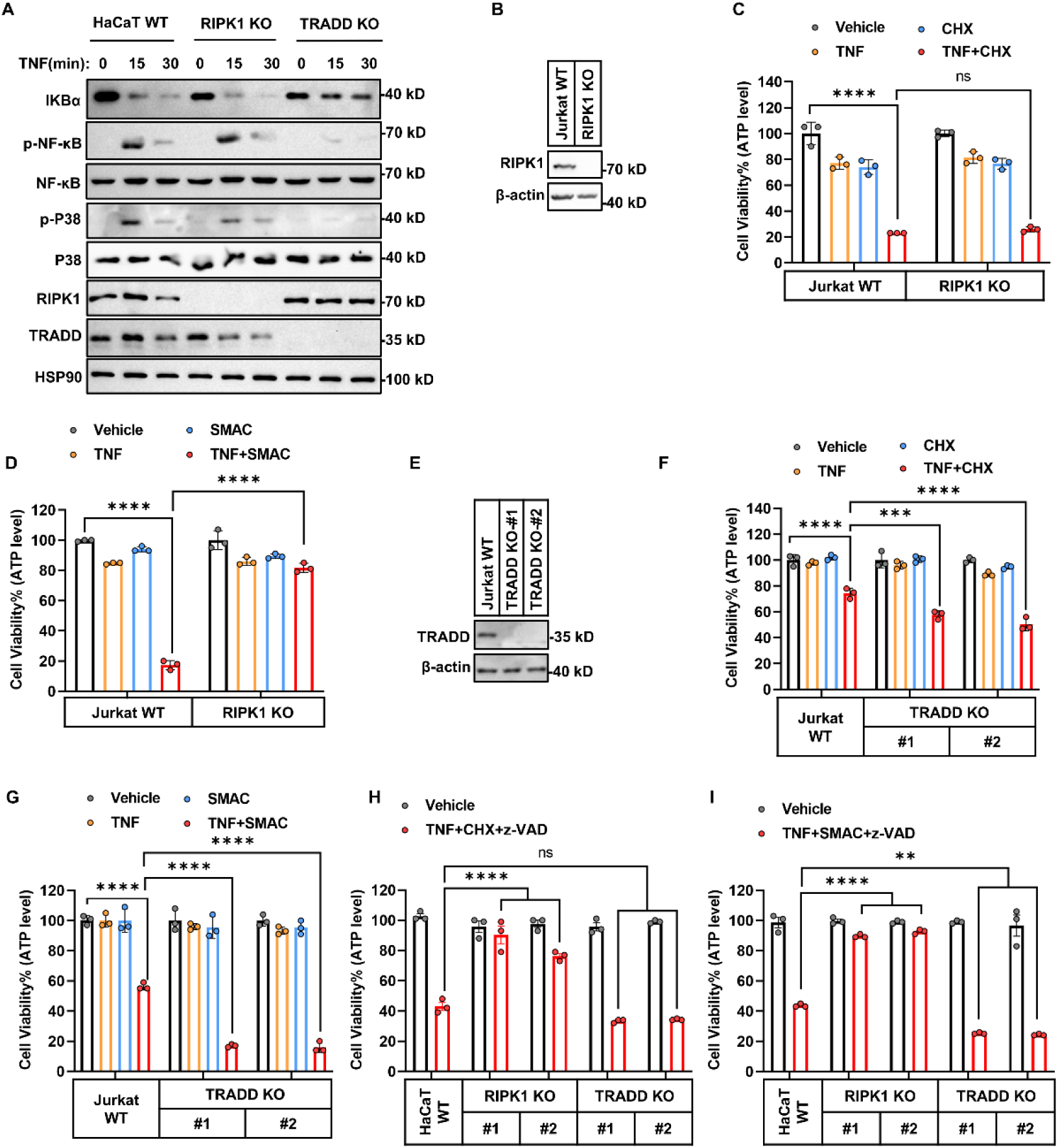
RIPK1 and TRADD are functionally redundant in kinase-independent apoptosis in human cells. (A) NF-κB pathway activation was analyzed by immunoblotting in WT, RIPK1 KO, and TRADD KO HaCaT cells treated with TNF for 0, 15, or 30 minutes (n = 3). (B-D) RIPK1 KO in Jurkat cells was confirmed by immunoblotting (B). Jurkat WT and RIPK1 KO cells were treated with TNF/CHX for 12 h (C), TNF/SMAC for 12 h (D), and cell viability was assessed using ATP-based assay (n = 3). (E-G) TRADD KO in Jurkat cells was confirmed by immunoblotting (E). Jurkat WT and TRADD KO cells (clones #1 and #2) were treated with TNF/CHX for 8h (G), TNF/SMAC for 8h (G), and cell viability was assessed via ATP-based assay (n = 3). (H, I) WT, RIPK1 KO, TRADD KO HaCaT cells (clones #1 and #2 from two different gRNAs) were treated with TNF/CHX/z-VAD for 8 h (H) or TNF/SMAC/z-VAD for 24 h (I), and cell viability was measured by ATP-based luminescence assay (n = 3). The data in all graphs are showed as mean ± SD. Representative immunoblots from three independent experiments are shown. Statistical significance was determined using two-way ANOVA for group comparisons (C, D, F, G-I): ****P < 0.0001; ***P < 0.001 (F, P=0.0007); **P < 0.01 (I, P=0.0059); ns, not significant.

**Figure S4.**
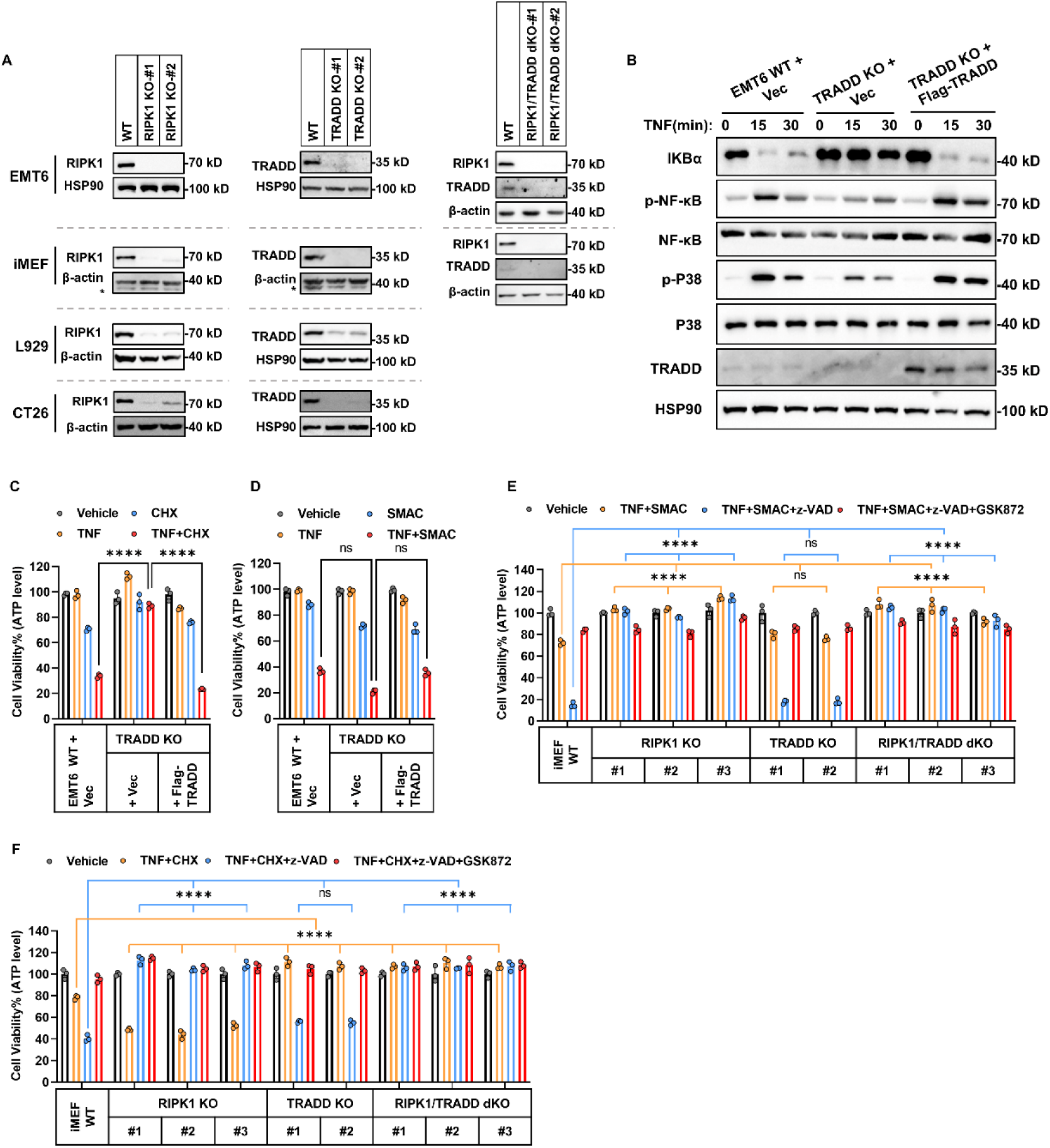
TRADD is indispensable for kinase-independent apoptosis in mouse cells. (A) RIPK1 KO, TRADD KO, RIPK1 and TRADD dKO in EMT6 or iMEF and RIPK1 KO and TRADD KO in L929 or CT26 cells (clones #1 and #2 from two different gRNAs) were confirmed by immunoblotting. (B) EMT6 WT, TRADD KO, and TRADD KO cells reconstituted with TRADD were treated with TNF for 0, 15, or 30 minutes. NF-κB and MAPK signaling activation was assessed by immunoblotting of key pathway proteins. (C-D) Vector-or TRADD-reconstituted EMT6 cells in both WT and TRADD KO backgrounds were treated with TNF/CHX for 8 h (C) or TNF/SMAC for 12 h (D). Cell viability was measured by ATP-based luminescence assay (n = 3). (E) iMEF cells (WT, RIPK1 KO clones #1–3, TRADD KO clones #1–2, RIPK1/TRADD dKO clones #1–3) were treated with TNF/SMAC, or TSZ in the presence or absence of RIPK3 inhibitor GSK872 for 12 h. Cell viability was measured using an ATP-based luminescence assay (n = 3). (F) The same panel of iMEF cells was treated with TNF/CHX or TCZ in the presence or absence of GSK872 for 6 h. Cell viability was measured using an ATP-based luminescence assay (n = 3). The data in all graphs are showed as mean ± SD. Representative immunoblots from three independent experiments are shown. Statistical significance was determined using two-way ANOVA for group comparisons (C-F): ****P < 0.0001; ns, not significant.

**Figure S5.**
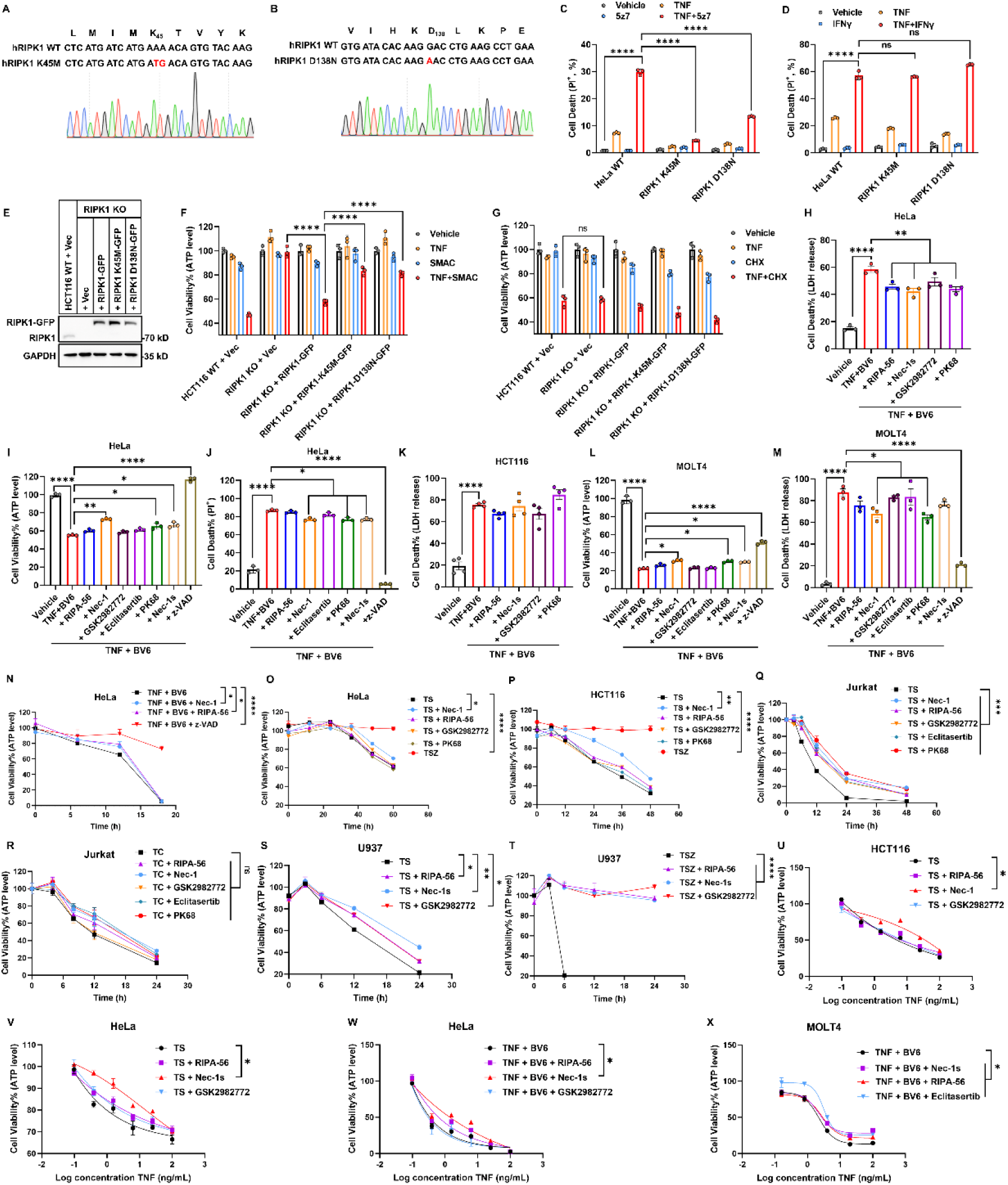
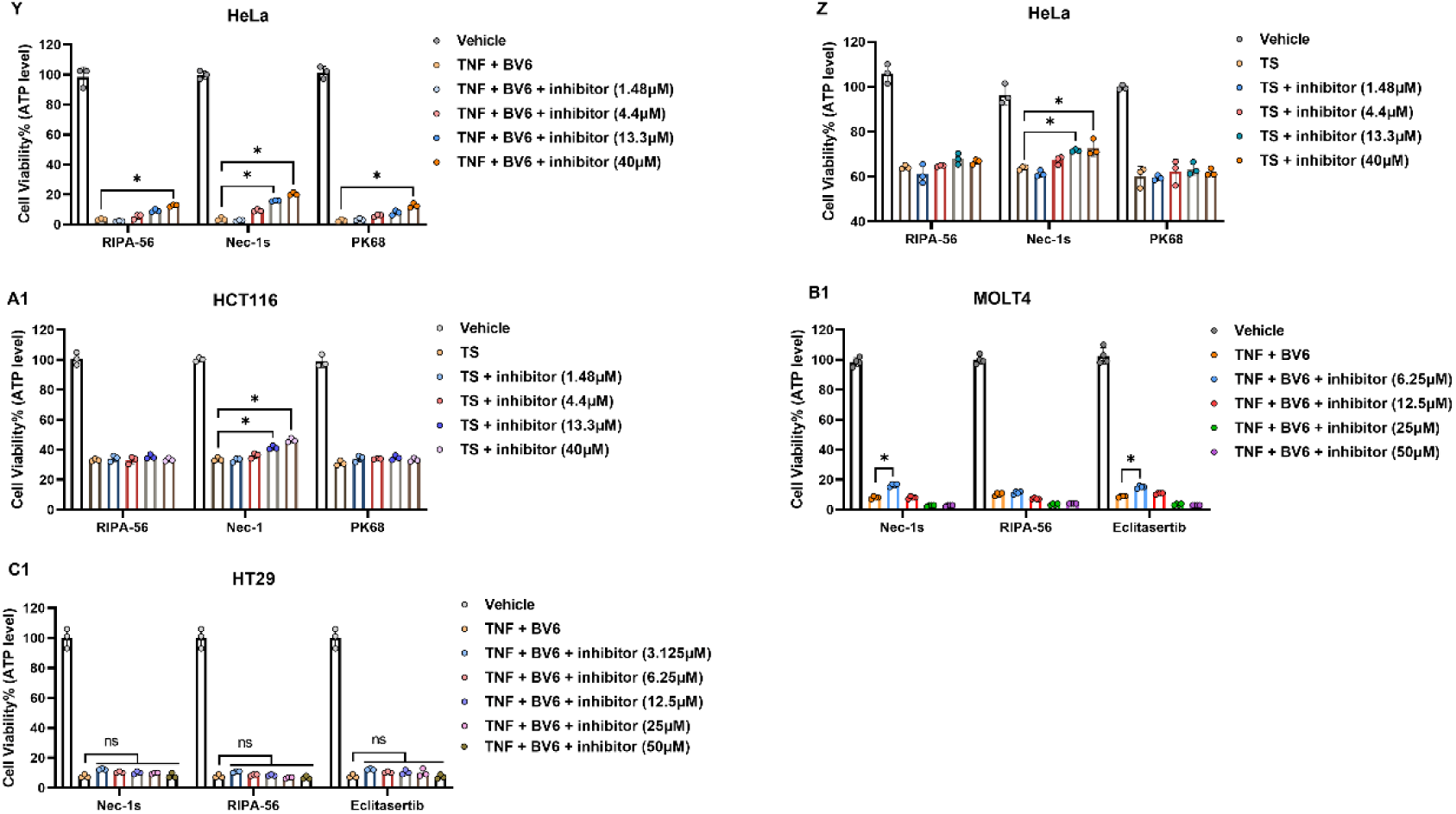
RIPK1 kinase inhibitors fail to block TNF/SMAC-induced apoptosis in various human cell types. (A, B) Sanger sequencing chromatograms confirming homozygous knock-in of RIPK1-K45M (A) and RIPK1-D138N (B) in HeLa cells. (C, D) HeLa WT, RIPK1-K45M, and RIPK1-D138N knock-in cells were treated with TNF/5Z7 for 48 h (C) or TNF/IFNγ for 48 h (D). Cell death was assessed by PI staining followed by flow cytometry (n = 3). (E-G) HCT116 WT, RIPK1 KO, and RIPK1 KO cells reconstituted with WT RIPK1-GFP, RIPK1-K45M-GFP, or RIPK1-D138N-GFP were analyzed for RIPK1 expression by immunoblotting (E). Cells were treated with TNF/SMAC for 24 h (F) or TNF/CHX for 8 h (G), and cell viability was measured by ATP-based luminescence assay (n = 3). (H-J) HeLa cells were treated with TNF/BV6 for 15 h, in the presence or absence of z-VAD and the indicated RIPK1 kinase inhibitors. Cell death was quantified by LDH release into the culture medium (H), an ATP-based luminescent viability assay (I), and PI staining followed by flow cytometry (J) (n = 3). (K) HCT116 cells were treated with TNF/BV6 for 12 h in the presence or absence of the indicated RIPK1 kinase inhibitors (RIPA-56, Nec-1s, GSK2982772, and PK68). Cell death was measured by LDH levels released in the culture medium (n = 4). (L, M) MOLT4 cells were treated with TNF/BV6 for 24 h, in the presence or absence of z-VAD and the indicated concentrations of RIPK1 kinase inhibitors (RIPA-56, Nec-1, GSK2982772, Eclitasertib, PK68, and Nec-1s). Cell viability was quantified by an ATP-based luminescent assay (L), and cell death was assessed by LDH release into the culture medium (M) (n = 3). (N, O) Viability of HeLa cells treated with TNF/BV6 (N), TNF/SMAC (O) in the presence or absence of z-VAD and the indicated RIPK1 inhibitors. Viability was measured by an ATP-based luminescence assay at different time points (n = 3). (P) Time-course viability of HCT116 cells treated with TNF/SMAC, in the presence or absence of z-VAD or RIPK1 inhibitors (n = 3). (Q, R) Time-course viability of Jurkat cells treated with TNF/SMAC (Q), TNF/CHX (R), in the presence or absence of RIPK1 inhibitors (n = 3). (S, T) Time-course viability of U937 cells treated with TNF/SMAC (S), TNF/SMAC/z-VAD (T), in the presence of indicated RIPK1 inhibitors (n = 3). (U) Dose-response viability of HCT116 cells treated with indicated concentrations of TNF and fixed SMAC, in the presence or absence of RIPK1 inhibitors. Cell viability was measured by ATP-based luminescence assay (n = 3). (V, W) Dose-response viability curves of HeLa cells treated with indicated concentrations of TNF with fixed SMAC (V), or BV6 (W), in the presence or absence of RIPK1 inhibitors (RIPA-56, Nec-1s, GSK2982772). Cell viability was measured by ATP-based luminescence assay (n = 3). (X) Dose-response viability of MOLT4 cells treated with indicated concentrations of TNF and fixed BV6, in the presence or absence of RIPK1 inhibitors. Cell viability was measured by ATP-based luminescence assay (n = 3). (Y) HeLa cells were treated with TNF/BV6, in the presence or absence of the indicated concentrations of RIPK1 inhibitors (RIPA-56, Nec-1s, PK68). Cell viability was measured by ATP-based luminescence assay (n = 3). (Z, A1) HeLa (Z) and HCT116 (A1) cells were treated with TNF/SMAC, in the presence of the indicated concentrations of RIPK1 inhibitors. Cell viability was measured (n = 3). (B1, C1) MOLT4 (B1) and HT29 (C1) cells were treated with TNF/BV6, in the presence or absence of RIPK1 inhibitors. Cell viability was measured via ATP-based luminescence assay (n = 3). The data in all graphs are showed as mean ± SD. Representative immunoblots from three independent experiments are shown. Statistical significance was determined using two-way ANOVA (C,D,F,G), or one-way ANOVA (H-C1) for group comparisons: ****P < 0.0001; ***P<0.001 (Q, P=0.0009);**P<0.01 (H, P=0.0085; I, P=0.0092; P, P=0.0085; S, P=0.0052); *P<0.05 (I, P=0.048, P=0.046, from left to right; J, P=0.047; L, P=0.036, P=0.038, P=0.045, from left to right; M, P=0.042; N, P=0.043, P=0.046, from left to right; O, P=0.041; S, P=0.025, P=0.032, from left to right; U, P=0.044; V, P=0.033; W, P=0.039; X, P=0.037; Y, P=0.041, P=0.039, P=0.035, P=0.043, from left to right; Z, P=0.047, P=0.043, from left to right; A1, P=0.045, P=0.041, from left to right; B1, P=0.042, P=0.043, from left to right); ns, not significant.

**Figure S6.**
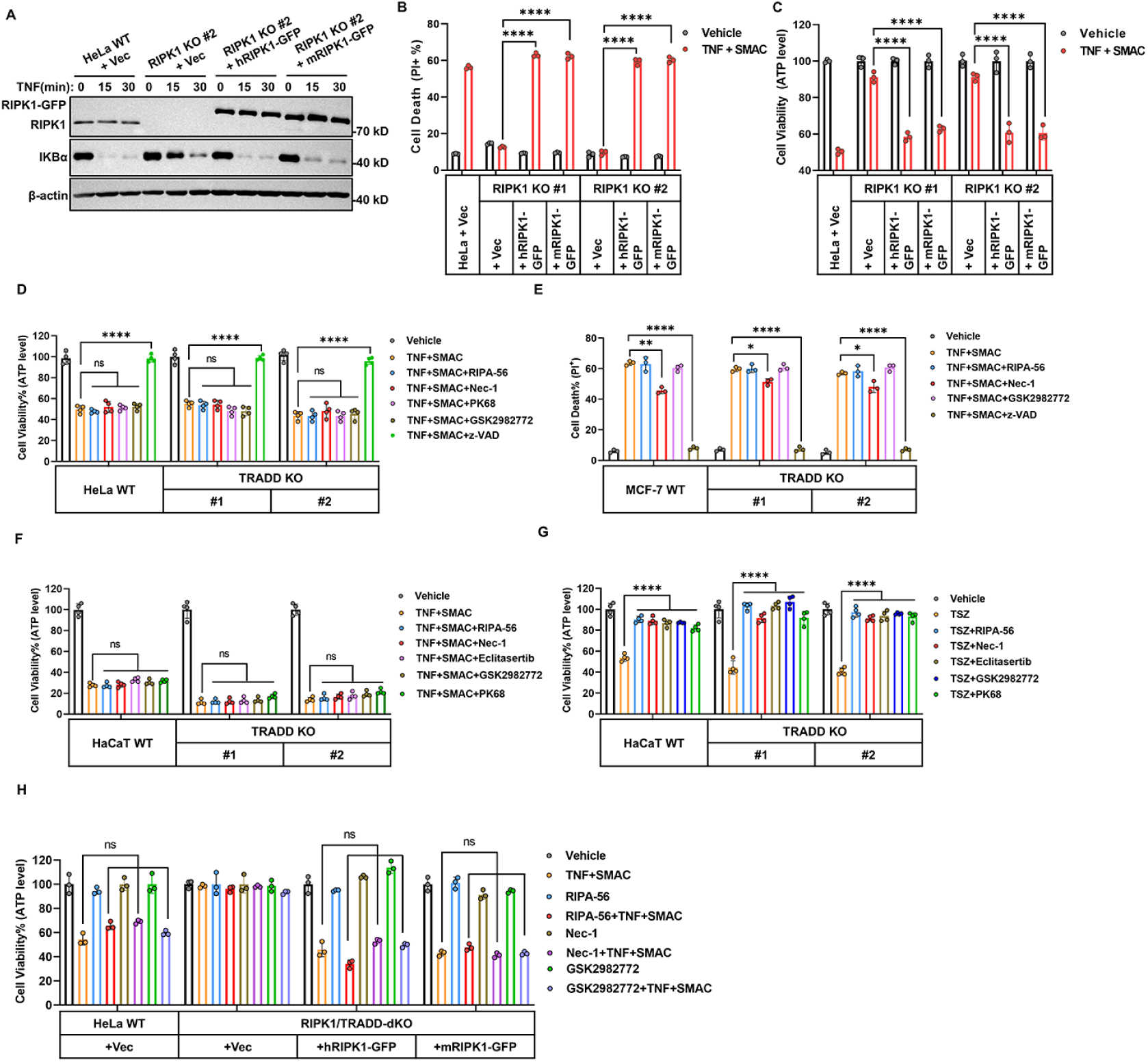
RIPK1 inhibitors fails to block TNF / SMAC induced apoptosis in human cells irrespective of RIPK1/TRADD status and remains ineffective after mouse RIPK1 reconstitution. (A) HeLa WT, RIPK1 KO clone #2, and RIPK1 KO clone #2 cells stably reconstituted with either human or mouse RIPK1-GFP were treated with TNF for 0, 15, or 30 min. Activation of NF-κB and MAPK signaling was assessed by immunoblotting (n = 3). (B, C) HeLa WT, RIPK1 KO clone #1 or #2, and RIPK1 KO clone #1 or #2 cells stably reconstituted with human or mouse RIPK1-GFP were treated with TS for 48 h. Cell death was evaluated by PI staining followed by flow cytometry (B), and cell viability was assessed by ATP-based luminescence assay (C) (n = 3). (D) HeLa WT or TRADD KO cells (clones #1 and #2) were treated with TNF/SMAC for 48h, in the presence or absence of RIPK1 kinase inhibitors (RIPA-56, Nec-1, PK68 or GSK2982772) and z-VAD. Cell viability was measured using an ATP-based luminescent assay (n = 4). (E) MCF-7 WT or TRADD KO cells (clones #1 and #2) were treated with TNF/SMAC for 48h in the presence of RIPK1 inhibitors or z-VAD. Cell death was measured by PI staining (n = 3). (F, G) HaCaT WT or TRADD KO cells (clones #1 and #2) were treated with either TNF/SMAC (F) or TNF/SMAC/z-VAD (G) for 24 h, in the presence of the indicated RIPK1 inhibitors. Cell viability was measured via an ATP-based assay (n = 4). (H)HeLa cells (WT, RIPK1/TRADD dKO, dKO reconstituted with human RIPK1-GFP or mouse RIPK1-GFP) were treated with TNF/SMAC in the presence of RIPK1 inhibitors (RIPA-56, Nec-1, or GSK2982772) for 48 h. Cell viability was measured (n = 3). The data in all graphs are showed as mean ± SD. Representative immunoblots from three independent experiments are shown. Statistical significance was determined using two-way ANOVA (B, C), or one-way ANOVA (D-H) for group comparisons: ****P < 0.0001; **P<0.01 (E, P=0.0051); *P<0.05 (E, P=0.035, P=0.025, from left to right); ns, not significant.

**Figure S7.**
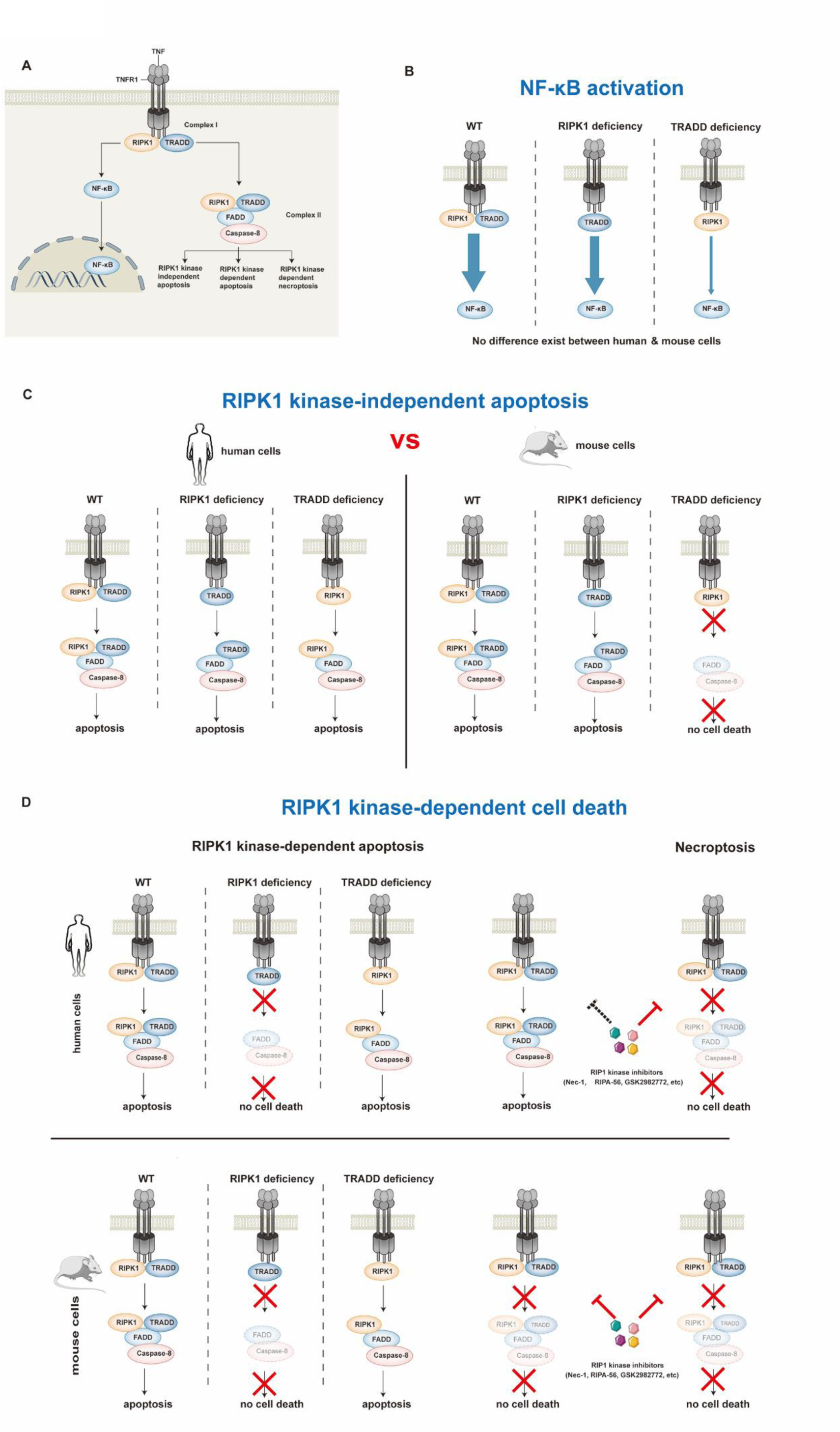
Differential roles of TRADD and RIPK1 in TNF-induced signaling and cell death pathways across species. (A) Schematic overview of TNFR1 signaling. Upon TNF stimulation, complex I forms at the plasma membrane containing TRADD and RIPK1, promoting NF-κB activation. Internalization leads to the assembly of cytosolic complex II, which mediates either RIPK1 kinase-independent apoptosis, RIPK1 kinase-dependent apoptosis, or necroptosis depending on the cellular context. (B) NF-κB activation is conserved across species and remains functional in both RIPK1-and TRADD-deficient cells. However, TRADD plays a more significant role than RIPK1 in supporting NF-κB signaling, as its deletion causes a stronger reduction in pathway activation. (C) Comparison of RIPK1 kinase-independent apoptosis in human and mouse cells. In human cells, RIPK1 and TRADD play redundant roles in mediating apoptosis; deletion of either does not prevent cell death. In contrast, TRADD is essential for RIPK1 kinase-independent apoptosis in mouse cells, as its deletion blocks the formation of the FADD–Caspase-8 complex and abolishes cell death. (D) Analysis of RIPK1 kinase-dependent apoptosis and necroptosis in human and mouse cells. In both species, RIPK1 is genetically essential for kinase-dependent cell death, whereas TRADD is dispensable. However, pharmacological inhibition of RIPK1 kinase activity shows species-specific effects: in mouse cells, RIPK1 kinase inhibitors (e.g., Nec-1, RIPA-56, GSK2982772) effectively block both RIPK1-dependent apoptosis and necroptosis; in human cells, these inhibitors only block necroptosis, but not RIPK1-dependent apoptosis.

**Supplementary Table 1.**
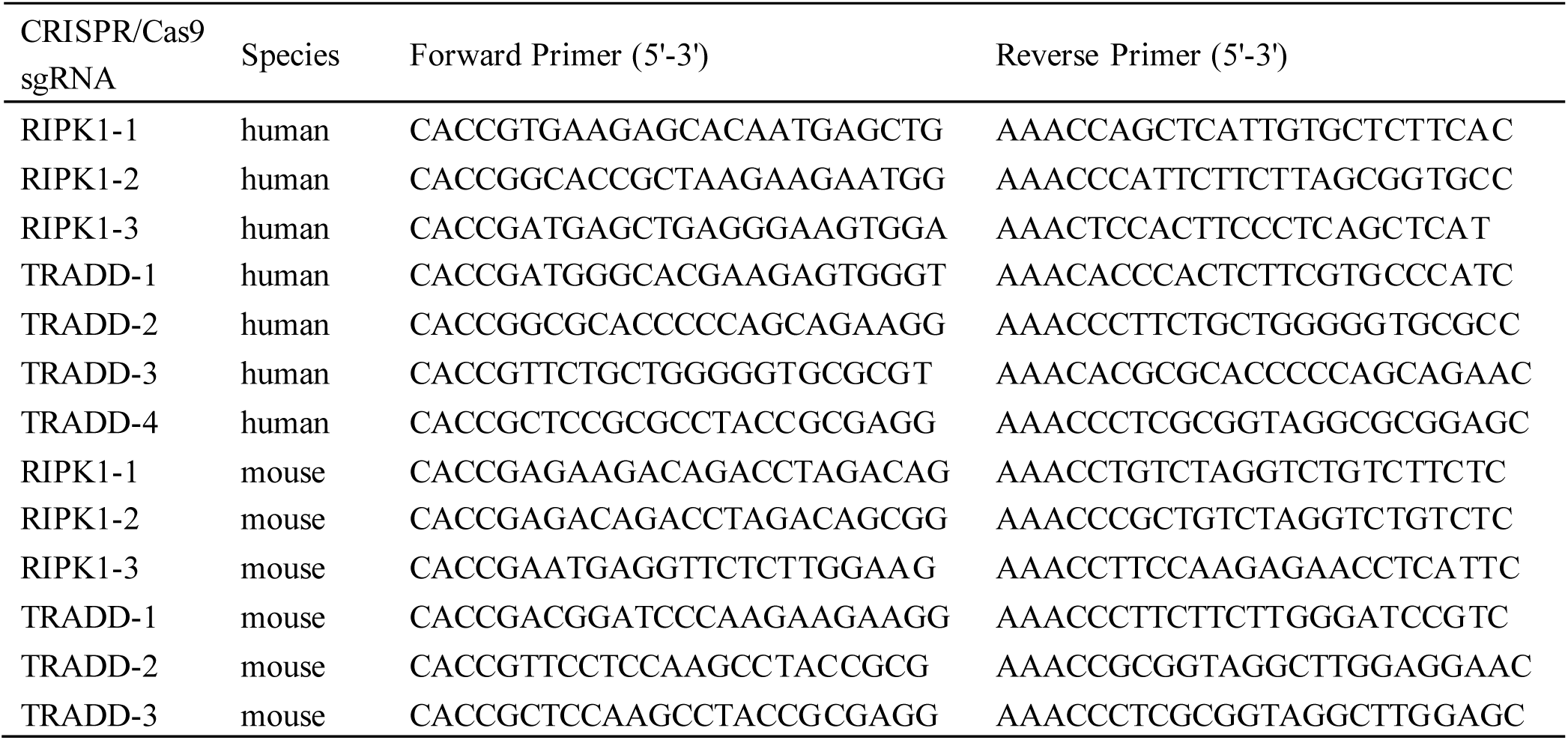
Primers used for sgRNA.

